# The structural variation landscape in 492 Atlantic salmon genomes

**DOI:** 10.1101/2020.05.16.099614

**Authors:** Alicia C. Bertolotti, Ryan M. Layer, Manu Kumar Gundappa, Michael D. Gallagher, Ege Pehlivanoglu, Torfinn Nome, Diego Robledo, Matthew P. Kent, Line L. Røsæg, Matilde M. Holen, Teshome D. Mulugeta, Thomas J. Ashton, Kjetil Hindar, Harald Sægrov, Bjørn Florø-Larsen, Jaakko Erkinaro, Craig R. Primmer, Louis Bernatchez, Samuel A.M. Martin, Ian A. Johnston, Simen R. Sandve, Sigbjørn Lien, Daniel J. Macqueen

## Abstract

Structural variants (SVs) are a major source of genetic and phenotypic variation, but remain challenging to accurately type and are hence poorly characterized in most species. We present an approach for reliable SV discovery in non-model species using whole genome sequencing and report 15,483 high-confidence SVs in 492 Atlantic salmon (*Salmo salar* L.) sampled from a broad phylogeographic distribution. These SVs recover population genetic structure with high resolution, include an active DNA transposon, widely affect functional features, and overlap more duplicated genes retained from an ancestral salmonid autotetraploidization event than expected. Changes in SV allele frequency between wild and farmed fish indicate polygenic selection on behavioural traits during domestication, targeting brain-expressed synaptic networks linked to neurological disorders in humans. This study offers novel insights into the role of SVs in genome evolution and the genetic architecture of domestication traits, along with resources supporting reliable SV discovery in non-model species.

## Main

Modern genetics remains primarily focused on single nucleotide polymorphism (SNP) analyses, with a growing recognition of the importance of larger structural variants (SVs) including inversions, insertions, deletions and copy number variations (CNVs) (defined here as variants ≥100 bp), among others^1^. SVs affect a larger proportion of bases in human genomes than SNPs^4^, are not always reliably tagged by SNPs^5^, more frequently have regulatory impacts^6^, and have been shown to alter the structure, presence, number, dosage, and regulation of many genes^1^. Nonetheless, SVs remain challenging to accurately type using whole genome sequence data^2-3^, limiting our understanding of their biological roles and exploitation as genetic markers. Consequently, there is a need for reliable SV detection approaches to fully exploit the fast-accumulating genome sequencing datasets in both model and non-model species, allowing for more complete genetics investigations. Many tools exist for SV discovery using short-read sequencing data, but all suffer from high false discovery rates (10-89%)^2,3,7^. This poses a challenge for truly *de novo* SV detection in previously unstudied species lacking ‘gold standard’ reference SVs to help distinguish true from false calls. Most studies rely on combining an ensemble of signals from different SV detection methods, although this strategy does not reliably improve performance and can in some cases aggravate false discovery^3^. Researchers therefore often apply independent experimental^8-9^ or visualization methods^10^ to validate a subset of SV calls. Overall, there remains an unsatisfactory lack of consensus on how to validate the quality of *de novo* SV datasets in most species^3^.

Salmonids have the highest combined economic, ecological and scientific importance among all fish lineages, and have consequently been subject to hundreds of genetics studies employing SNPs and other molecular markers^11,12^. In common with most non-model fish species, the SV landscape remains extremely poorly characterized in salmonids, apart from recent work informed by SNPs that revealed multi-megabase inversions in rainbow trout (*Oncorhynchus mykiss* Walbaum) influencing migration^13,14^, and a chromosomal fusion under selection in Atlantic salmon^15^, consistent with roles in adaptation. Salmonids offer a unique system to characterize SVs due to an ancestral salmonid-specific autotetraploidization (i.e. whole genome duplication, WGD) event (Ss4R), which occurred 80-100 Mya, following an earlier WGD (300-350 Mya) in the teleost common ancestor^16,17,18^. WGD events may influence selection on SV retention due to the functional redundancy linked to mass retention of duplicated genes, though this idea is yet to be tested. In addition, salmonids have been farmed in aquaculture for a small number (<15) of generations^11^, and while the genetic architecture of such recent domestication has been investigated using SNPs^19^, the role played by SVs remains unexplored. Finally, the application of SVs in selective breeding of salmonids and other commercial fishes remains untested. Clearly, the lack of SV data and analysis frameworks in salmonids represents an important knowledge gap.

Here we provide an end-to-end workflow to detect, genotype, validate and annotate SVs using short-read sequencing, removing false positives through efficient manual curation^10^, allowing reliable SV discovery in non-model species. Using this approach, we report a detailed investigation of the genomic landscape of SVs in the iconic Atlantic salmon, inclusive of 492 genomes representing wild and farmed genetic diversity, and populations of both European and North American descent.

## Results

### Accurate SV discovery in Atlantic salmon

We developed a workflow for SV discovery using paired-end short-read sequencing data aligned to the unmasked ICSASG_V2 reference assembly^17^, which can be run in Snakemake^20^ (Supplementary Figure 1). The probabilistic tool Lumpy^21^ was used for SV detection, which simultaneously draws on multiple evidence and SVtyper^22^ was used for genotyping. As *de novo* SV detection using short-read data is prone to false positives^3,21,23^, we added steps to avoid SV calling in complex regions of the genome where false positive rates were predicted to be particularly high (proven below). This included regions of ≥100x coverage (>10 times higher than the global average of 8.1x coverage across 492 samples), shown elsewhere to be overwhelmingly false calls^3^, as well as gap regions in the ICSASG_V2 assembly. These complex regions were most prevalent in chromosome arms where rediploidization was delayed after Ss4R, characterized by high sequence similarity among duplicated regions^17^ (Supplementary Figure 2).

Rather than using evidence from additional SV detection tools as a filter for true SV calls, a strategy shown elsewhere to be potentially unreliable^3^, we applied a curation approach to the entire filtered SV dataset using SV-plaudit^10^. SV-plaudit is a scalable framework for the rapid production of thousands of SV images via Amazon web services^10^ (examples: Supplementary Figures 3-8). This approach allowed us to efficiently retain high-confidence SV calls, while excluding low confidence or ambiguous calls, on the basis of available visual evidence drawn from paired-end and split-read alignments, in addition to read depth^10,21^. The Atlantic salmon individuals (Supplementary Table l) produced on average 55,754 SV calls (median: 55,041, SD: 10,051) before filtering complex regions and SV-plaudit curation (Supplementary Table 2). Across all individuals, 165,116 unique SVs were detected (size: 100bp to 2 million bp), which included an outlier peak of deletion SVs in the 1,432-1,436 bp size range (Supplementary Data 1; Supplementary Figure 9).

Using SV-plaudit on the full set of SV calls allowed us to retain only high-confidence calls, quantify the impact of filtering complex regions, and estimate a false discovery rate (FDR). The overall estimated FDR was 0.91 (149,491/165,116 of calls had low confidence), in line with the highest estimates in the literature^2,3,7^. In complex regions, the FDR was 0.992 (47,268/47,636 calls had low confidence). In the remaining chromosome-anchored assembly, the FDR was 0.85, validating the usefulness of removing complex genomic regions. Sequencing depth was not a reliable indicator of FDR (Supplementary Figure 10). A final high-quality set of 15,483 unique SV calls (14,017 deletions, 1,244 duplications, 242 inversions) and their genomic location is visualized in Fig. 1a and 1b. The average size for deletions was 1,532 bp (100 to 1,946,935 bp; SD: 23,070 bp) and for duplications 8,183 bp (102 to 80,1673 bp; SD: 25,589 bp) (Fig. 1c, d). For inversions, the average size was 121,935 bp (113 to 1,796,230 bp; SD: 278,698 bp) (Fig. 1e). The outlier peak at 1,432-1436 bp remained in the high-confidence deletions (Fig. 1c).

**Fig. 1.**
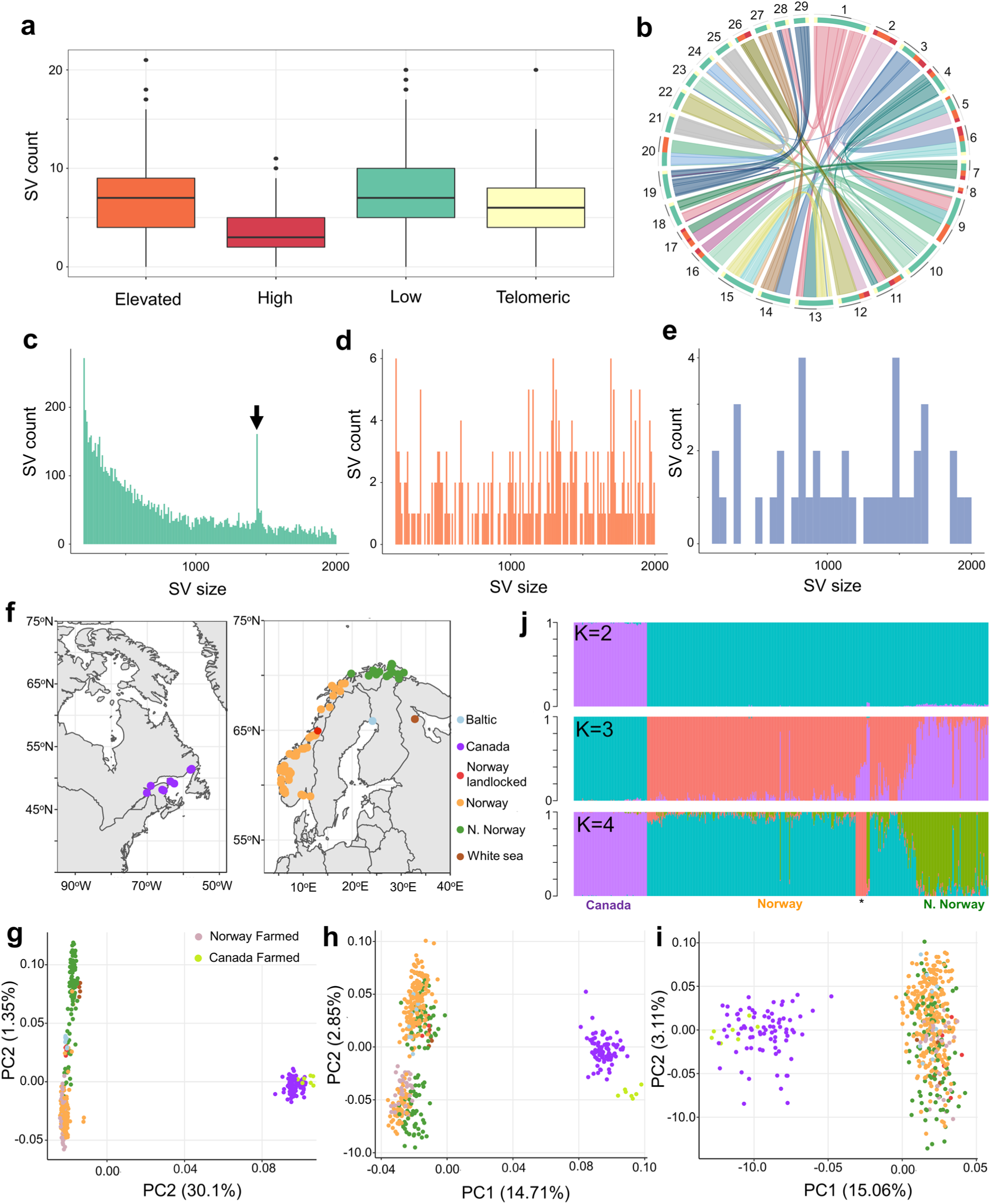
SV landscape in 492 Atlantic salmon genomes. **a** SV counts per 1-million bp window in the genome split into homology categories^17^ representing duplicated regions retained from the Ss4R WGD sharing ‘low’ (<90% identity), ‘elevated’ (90-95% identity) and ‘high’ (>95% identity) similarity in addition to telomere regions. **b** Locations of the same regions depicted on a Circos plot using the same colour scheme. **c-e** Size distributions of SVs for deletions (**c**), duplications (**d**) and inversions (**e**) with X axis limited to SVs ≤ 2,000 bp. Arrow in part **c** marks outlier peak in deletion calls (see Fig. 2). **f** Sampling locations of wild populations. **i-h** PCA of for each SV class: 14,017 deletions (**g**), 1,244 duplications (**h**), 242 inversions (**i**) with population matched by colour to part **f** for wild fish, and additional symbols given for farmed fish. **j** NGSadmix^86^ analysis of 14,017 deletions with K=2, 3 and 4. Each individual is a vertical line with different colours marking genetically distinct groups. Asterisk corresponds to White sea, Baltic and landlocked populations (K=4 plot).

To validate our SV discovery workflow we estimated the true positive rate for SV presence/absence and genotype calls using the high-confidence data retained after the SV-plaudit step. We sequenced PCR amplicons for 876 independent SV calls representing 168 unique SVs (108 deletions, 46 duplications, 15 inversions) (Supplementary Figure 11) at ≥50x coverage on the MinION platform. Across all SV calls, the true positive rate was 0.88 for SV presence/absence and 0.81 for SV plus genotype. For deletion calls, the true positive rate was 0.93 for presence/absence (520/559 calls) and 0.85 (475/559 calls) for genotype. For duplications, the true positive rate was 0.81 for presence/absence (186/230 calls) and 0.74 (170/230 calls) for genotype. For inversion calls, the true positive rate was 0.78 for presence/absence (68/87 calls) and 0.75(65/87 calls) for genotype. Full results are shown in Supplementary Table 3 (with examples in Supplementary Figures 12, 13 and 14). In summary, SV-plaudit curation vastly reduced the FDR to maintain predominantly true SV calls (provided in Supplementary Data 2).

To further confirm data quality, we asked if the high confidence SVs genotypes capture expected population genetic structure (Fig. 1f-j). SV genotypes were used in principal component analyses (PCA) for the different SV types (Fig. 1f-i). For all SV types, PC1 separated European and Canadian salmon, consistent with past work e.g.^24,25^. Deletions achieved a better resolution for the sampled European populations, with PC2 separating populations from Europe into distinct groups explained by latitude with evidence of intermixing at middle latitudes in Norway (Supplementary Figure 15), as reported elsewhere^24^. All farmed salmon clustered with the wild populations from which they are descended. Farmed salmon from Europe, including 13 farmed fish from Chile, clustered with wild salmon from Southern Norway, while 7 Chilean farmed salmon clustered with Canadian salmon (Fig. 1c). Using the high-confidence deletion genotypes, an admixture analysis was performed, which was consistent with the PC analysis (Fig. 1j). For comparison, we also performed PCAs using the raw unfiltered SV calls, plus the reduced subset filtered for complex regions, which failed to capture the same population structure (Supplementary Figure 16). In summary, our final set of deletion genotypes capture expected population genetic structure at the highest resolution. It is unclear if the weaker signal for duplications and inversions is linked to specific properties of these markers, their comparatively lower number, or slightly lower genotyping accuracy.

### Annotation of Atlantic salmon SVs

We used SnpEff^26^ to annotate all high confidence SV calls against features in the ICSASG_v2 annotation. Many SVs were located in intergenic and intronic regions (Supplementary Figure 17), with 62%, 3% and 2.5% within 5 kb of a protein-coding gene, long non-coding RNA gene or pseudogene, respectively. Around half (49%) of all SVs overlapped one or more RefSeq gene, the majority of which overlapped a single gene (Supplementary Figure 18), with 8,439 genes overlapped in total. Approximately 4%, 21% and 25% of deletions, duplications and inversions were predicted by SnpEff to have a high impact, respectively, including hundreds of putative exon losses, frameshift variants and potential gene fusion events (Supplementary Figure 19). 101 duplications spanned entire genes (mean length: 51.7 kb, median length: 15.1 kb). The high impact annotations for different SV types were associated with an overrepresentation of several biological processes in the gene ontology (GO) framework^27^ (Supplementary Table 4, 5).

### Recently active DNA transposon in *Salmo* evolution

The outlier peak observed in the deletion calls (Fig. 1a; Supplementary Figure 9) was investigated by extracting all high confidence variants of 1,432-1,436 bp in size (104 sequences) from the ICSASG_v2 genome. 94 and 89 of these sequences shared ≥50% and ≥95% identity in all pairwise combinations, respectively. The 94 sequences were used as queries in BLASTn searches revealing that 91% (86 out of 94) shared ≥95% identity to a pTSsa2 piggyBac-like DNA transposon (NCBI accession: EF685967)^28^. The breakpoints in the outlier deletions SV match to the complete pTSsa2 sequence (Supplementary Data 3), missing no more than a few bp at the 5’ or 3’ end. Consequently, the outlier deletion peak (Fig. 1a) appears to largely represent an intact pTSsa2 sequence.

Phylogenetic analysis was done incorporating the Atlantic salmon pTSsa2 sequences along with the top 100 BLASTn hits to the pTSsa2 sequence in the genome of brown trout *Salmo trutta* (repeat masking off; all sequences e-value = 0.0, 70-100% and 84-95% query coverage and identity, respectively). Repeating the search against genomes for the next most closely-related salmonid genera, *Salvelinus* (Arctic charr *S. alpinus*) and *Oncorhynchus* (rainbow trout *O. mykiss*, coho salmon *O. kitsuch*, and chinook salmon *O. tshawytscha*) failed to identify sequences sharing >50% coverage or >81% identity. The tree indicates independent expansions of pTSsa2 sequences in the Atlantic salmon and brown trout genome (Fig. 2_;_ Supplementary Figure 20). The pTSsa2 sequence appears in the Atlantic salmon genome with high copy number across all chromosomes (Supplementary Figure 21).

**Fig. 2.**
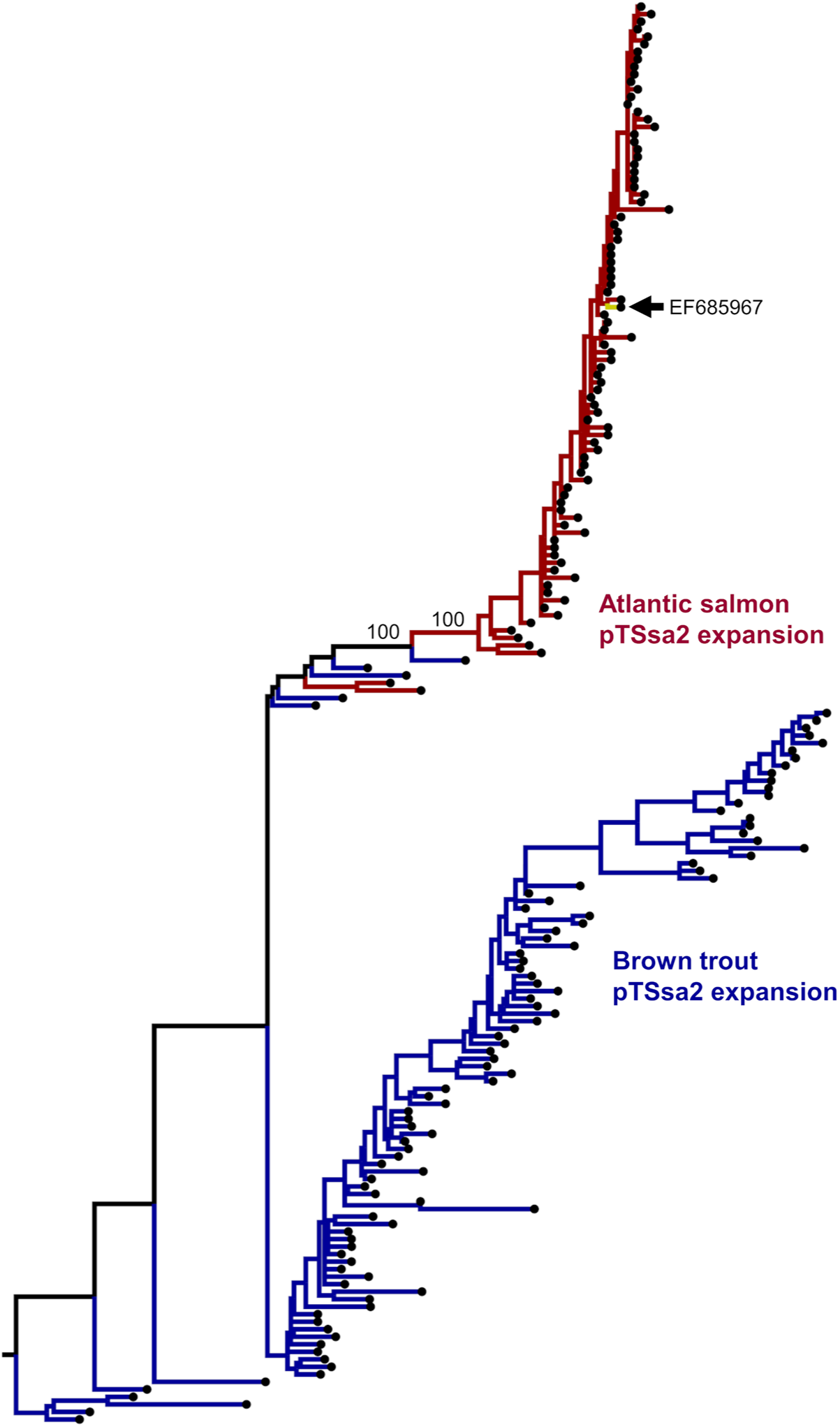
Evidence for an active DNA transposon in *Salmo* evolution. Phylogenetic tree of Atlantic salmon sequences representing deletion polymorphisms matching the pTSsa2 piggyBac-like DNA transposon^28^ (EF685967) and 100 top hits to this sequence within the brown trout genome. The tree was generated from an alignment spanning the length of pTSsa2 (Supplementary Data 3) using the TPM3+F+G4 substitution model. Bootstrap values are given at key nodes. A full tree with sequence identifiers, genomic locations of pTSsa2 sequences and bootstrap values is provided in Supplementary Fig. 18. A circos plot highlighting the location of pTSsa2 sequences in the Atlantic salmon genome is given in Supplementary Fig. 19.

We also determined the broader overlap of SVs and repeat sequences in the Atlantic salmon genome. Among all SVs, 65% (10,184) contained no repeat sequences, 16% (2,423) a single repeat, and 7% (1,027) two repeats. There was a significant correlation between SV size and the number of repeats per SV across all SV types (Pearson’s R ≥0.99, *P* < 0.0001 in each test), indicating that the number of repeats within each SV was simply a direct product of SV size.

### Impact of genome duplication on the SV landscape

Salmonid genomes retain a global signature of duplication from Ss4R, with at least half of the protein-coding genes retained as expressed, functional duplicates (referred to as ohnologs)^17,18^. Ss4R ohnolog pairs share amino acid sequence identity ranging from ∼75 to 100%^12,17,18^ with ∼40% maintaining the ancestral tissue expression pattern^17^, suggesting pervasive functional redundancy. We hypothesised that the redundancy provided ohnolog retention after WGD influenced the evolution of the SV landscape by creating a mutational buffer^29^ against deleterious SV mutations. A key prediction is that genes found in Ss4R ohnolog pairs (with scope for functional redundancy) should be more overlapped by SVs compared to singleton genes (lacking scope for functional redundancy).

We tested this prediction by generating a novel set of high-confidence Ss4R ohnolog pairs (10,023 pairs, i.e. 20,046 genes) and singletons (8,282 genes) (Supplementary Data 4) and indeed, found a significant enrichment of SVs overlapping retained Ss4R ohnologs (*Fisher’s exact test, P* = 0, odds ratio = 1.47) (Supplementary Table 6). This effect was specific to deletions (*P* = 0, odds ratio = 1.62), and hence not observed in duplications (*P* = 0.62) nor inversions (*P* = 0.52). SVs with putative high impact did not overlap ohnologs more than singletons (high impact snpEff annotation: *P =* 0.93, manually curated deletions impacting exons: *P =* 0.55) (Supplementary Data 5).

Next we asked if gene expression characteristics influence the overlap between SVs and Ss4R ohnologs. We initially used Spearman’s rank correlation to establish co-expression of ohnologs across an RNA-Seq atlas of 15 tissues^16^. We found that ohnolog pairs where one copy overlaps an SV showed slightly lower expression correlation compared to randomly selected ohnolog pairs (resampling test, *P* = 0)

(Supplementary Figure 22). This pattern could be explained by SVs affecting ohnolog pairs with greater levels of functional divergence, but may also be caused by relaxed purifying selection on duplicated copies, allowing more SVs to accumulate. It has been shown elsewhere that the more highly expressed ohnolog in a pair is typically under stronger purifying selection^30^. Therefore, we asked if ohnologs overlapped by a deletion SV have reduced expression compared to their duplicate with no SV overlap. Indeed, this was the case (*Wilcoxon rank-sum test, P* = 2.9e-6) (Supplementary Figure 22). We also found that ohnolog pairs showing overlap with deletion SVs showed reduced expression compared to ohnolog pairs showing no overlap to SVs (*Wilcoxon rank-sum* test, *P* = 7e-25) (Supplementary Figure 22).

Overall, these analyses reveal that the Ss4R WGD strongly influenced the retention of deletion SVs in the Atlantic salmon genome, and this may be explained by functional redundancy.

### Selection on SVs during Atlantic salmon domestication

Our study provides a unique opportunity to ask if SVs were selected during the domestication of Atlantic salmon, which commenced when the Norwegian aquaculture industry was founded in the late 1960s^11,31^. Consequently, farmed Atlantic salmon are no more than 15 generations ‘from the wild’, in contrast to livestock and poultry, which have been domesticated for thousands of years^11,12^. The early domestication process involves strong selection on behavioural traits^32,33^ targeting molecular pathways underpinning cognition, learning and memory, for instance genes with functions in synaptic transmission and plasticity^34,35^. Specifically, selection on farmed animals should remove individuals that invest in costly behavioural and stress responses such as predator avoidance and fear processing, in favour of animals that invest into performance traits ^32,36^. We thus hypothesised that SVs linked to genes regulating pathways controlling behaviour would be under distinct selective pressures in farmed and wild salmon.

To test our hypothesis, we established significantly genetically differentiated SVs by calculating the fixation index (F_ST_)^37^ between 34 farmed Norwegian salmon and 257 wild salmon from Norway. The wild individuals were selected based on a PCA including all European salmon, aiming to remove confounding effects of genetic differentiation by latitude observed in wild Norwegian salmon (Fig. 3a), retaining the closest possible background to the wild founders used in aquaculture. We used a permutation approach to estimate the probability of observed F_ST_ values in relation to random expectations, defining 584 SV outliers at *P*<0.01 (all F_ST_ >0.103, Median F_ST_ = 0.149) (Fig. 3b; Supplementary Data 6), which were distributed throughout the genome (Fig. 3c).

**Fig. 3.**
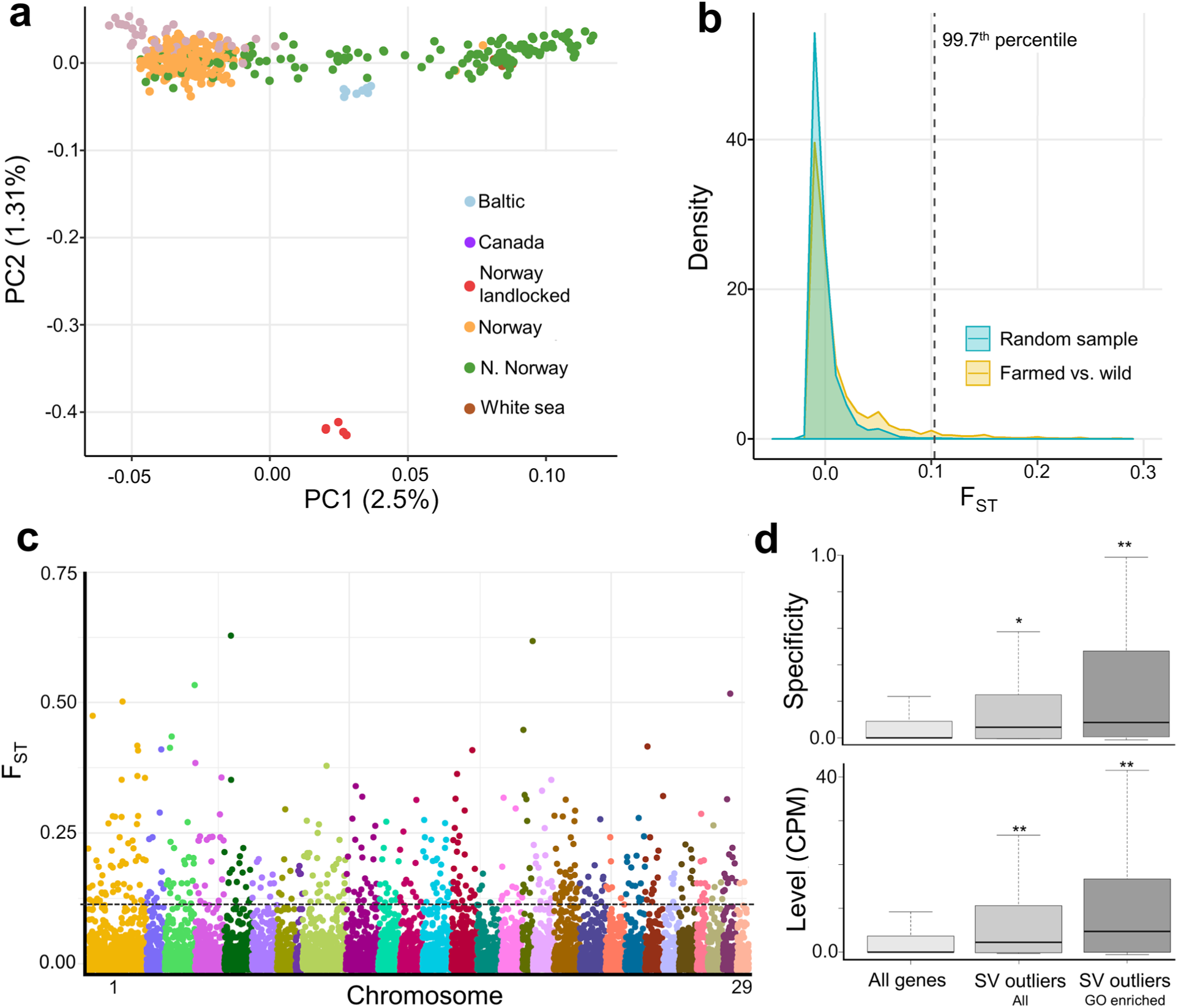
Genetic differentiation of SVs between farmed and wild Atlantic salmon. **a** PCA used to select appropriate wild individuals for F_ST_ comparison (n=257) vs. farmed salmon (n=34) on the basis of genetic distance by latitude (see also Supplementary Fig. 15) separated along PC1. The population symbols are the same as shown in Fig. 1. **b** Observed F_ST_ value distribution comparing farmed vs. wild salmon contrasted against 200 random distributions for the same number of individuals. Dotted line shows cut-off F_ST_ value employed in addition to a per SV criteria of *P*<0.01. **c** Manhattan plot of 12,627 F_ST_ values with dotted line showing the same cut-off above which are the 584 SV outliers. **d** Brain gene expression specificity (top panel) and expression level (bottom panel) are increased for genes linked to the 584 outlier SVs, with the effect more pronounced for a 326 gene subset contributing to significantly enriched GO terms, compared to 44,469 genes in a multi-tissue transcriptome. Single and double asterisks indicate *P<*0.005 and *P<*0.00001, respectively. The observed increase in expression was specific to brain (plots for other tissues shown in Supplementary Fig. 22 and 23). Statistical analysis for all tissues shown in Supplementary Table 9.

GO enrichment tests identified 132 overrepresented biological processes (*P*<0.05) among the genes linked to these outlier SVs by SnpEff (Supplementary Table 7). This set comprises 326 unique genes contributing to the enriched terms (Supplementary Table 8). 34 biological processes explained by 156 unique genes (48% of the unique genes contributing to all enriched GO terms) were daughter terms related either to learning and behaviour, including ‘habituation’ (*P*<0.002), ‘vocal learning’ (*P*<0.001), and ‘adult behavior’ (*P*<0.02), or the nervous system, including ‘positive regulation of nervous system process’ (*P*<0.02),’ presynaptic membrane assembly’(*P*<0.01), ‘postsynapse assembly’ (*P*<0.02) ‘oligodendrocyte development’ (*P*<0.001) and ‘regulation of neuronal synaptic plasticity’ (*P*<0.03).

To test our hypothesis, we asked if genes linked to outlier SVs showed enrichment in brain expression (Fig. 3d). Indeed, this was strongly supported when judged against transcriptome-wide expectations (Fig. 3d); with the signal being strongest for the 326 gene subset contributing to the overrepresented GO terms, emphasising particular importance of brain functions among the enriched gene set (Fig. 3d, Supplementary Table 9). A positive enrichment in the expression of outlier linked genes was only observed in brain, with nine other tested tissues showing either no differences to transcriptomic expectations, or in the case of muscle and foregut, reduced expression specificity (Supplementary Table 9; Supplementary Figures 23, 24). Finally, we asked if the outlier SVs overlapped putative cis-regulatory elements (CREs) detected in brain using novel ATAC-Seq data (significant peaks overlapping a gene +/- 3,000bp up/downstream; n=4) more than expected. For 9,920 SVs lacking evidence for differentiation between farmed and wild fish (F_ST_ *P* >0.05), 7.1% overlapped at least one brain ATAC-Seq peak, which was almost identical to SV outliers (7.0%) (*Fisher’s exact test, P* = 0.86). A similar result was observed by restricting the analysis to genes with brain biased expression (*Fisher’s exact test, P* = 0.41).

### SVs selected by domestication are linked to many synaptic genes

The increased brain expression and overrepresentation of nervous system functions for SV outlier linked genes motivated us to investigate the role of these loci in the genetic architecture of domestication. We performed a detailed annotation of the 156 SV outlier linked genes contributing to the 34 aforementioned enriched GO terms (Supplementary Table 10). To cement the relevance of this gene set to our hypothesis, we cross-referenced all the encoded protein products with a high-resolution synaptic proteome from zebrafish^38^. Our rationale was that the synaptic proteome is central to nervous system activity and defines the repertoire of cognitive and behaviours an animal can perform during its life^38,39^.

Among the 156 SV outlier linked genes, 65 (i.e. 42%, linked to 67 distinct SVs) encode a protein with an ortholog in the zebrafish synaptic proteome (Supplementary Table 10) defined by stringent reciprocal BLAST (mean respective pairwise % identity and coverage = 77% and 95%). As synaptic proteomes are highly conserved between fish and mammals^38^, it is reasonable to assume these proteins are *bone fide* components of Atlantic salmon synaptic proteomes, and that a minimum of 11% of the outlier SVs were linked to synaptic genes by SnpEff. These proteins are encoded by multiple members of ancient, conserved gene families involved in synaptic formation, transmission and plasticity, including neurexins (*NRXN1* and *NRXN2*), SH3 and multiple ankyrin repeat domains 3 proteins (*SHANK2* and *3*), cadherins (*CDH4, CDH8, CDH11, PCDH1*), Down syndrome cell adhesion molecules (*DSCAM* and *DSCAML*), teneurins (*TENM1* and *TENM2*), gamma-aminobutyric acid receptors (*GABRB2* and *GABRG2*), potassium voltage-gated channel subfamily D members (*KCND1* and *KCND2*), receptor-type tyrosine-protein phosphatases (*PTPRG* and *PTPRN2*) and ionotropic glutamate receptors (*GRIK3* and *GRIN2C*) (Fig. 4). Genetic disruption to orthologs for most these proteins (59/65) cause behavioural and/or neurological disorders in mammals (Supplementary Table 10).

**Fig. 4.**
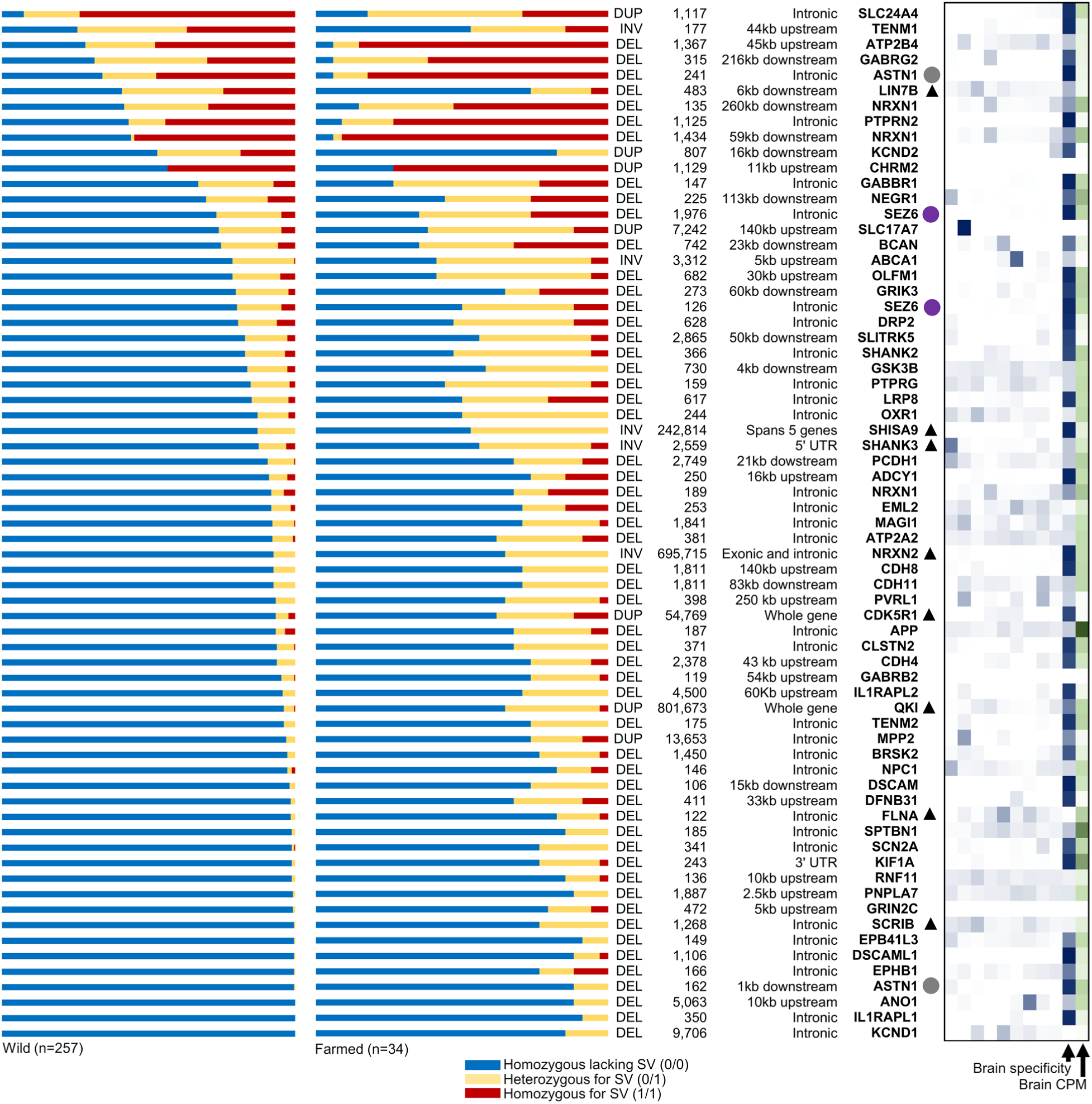
SVs under selection during Atlantic salmon domestication are linked to 65 unique genes encoding synaptic proteins. SV genotypes are visualized on the left, ordered from bottom to top with decreasing frequency of homozygous genotypes (0/0) lacking the SV in wild fish. Annotation of each SV type, its size, and genomic location with respect to each synaptic gene is also shown. The circles next to genes highlight Ss4R ohnolog pairs and the black triangles indicate the overlap of an SV with a putative cis regulatory element (ATAC-Seq peak). The heatmap on the right depicts the expression specificity of each gene across an RNA-Seq tissue panel^17^ (white to dark blue depicts lowest to highest tissue specificity; tissues shown in different columns from left to right: liver, gill, skeletal muscle, spleen, heart, foregut, pyloric caeca, pancreas, brain). The overall expression of each gene in brain is shown on the right of the heatmap (white to dark green depicts increasing CPM across the column). Data provided in Supplementary Table 10.

To ask how selection acted on these variants during domestication, we compared allele frequencies between wild and farmed fish (Fig. 4). By far the most common scenario was that the synapse gene - linked SVs are rare alleles in wild fish that show increased frequency of heterozygotes (carrying one SV copy, 0/1) and homozygotes (carrying both SV copies, 1/1) in farmed fish (Fig. 4). We also found that farmed individuals often carry multiple copies of SVs that are especially rare in wild fish (defined as 0/0 homozygous frequency ≥ 0.90, 45 SVs) - assumed to be deleterious in natural environments - including homozygote 1/1 states for SVs located on different chromosomes (Supplementary Figure 25).

Many of the outlier SVs linked to the 65 synaptic genes are located in non-coding regions (introns and untranslated regions, 45%), while a smaller fraction are located within 10kb up or downstream (15%) or within ≥10kb to 260 kb (33%) of the same genes (Fig. 4). A smaller fraction affect coding regions via whole gene duplications, either involving a small number of genes, e.g. a 55 kb duplication overlapping the brain-specific *CDK5R1* gene, or through larger multigene duplications (Fig. 4; Supplementary Table 10). A striking example of an SV with a putative major disruptive effect was a 696 kb inversion that flips multiple exons and the upstream region of the brain-specific gene encoding neurexin 2, which should halt translation of a functional protein (Supplementary Table 10). Finally, among this synaptic gene set, we identified two ohnolog pairs retained from Ss4R encoding astrotactin-1 and seizure protein 6 (Fig. 4).

### Major effect SVs altered by domestication

We identified 32 further SVs with major predicted effects on gene structure and function among the significant F_ST_ outliers, which typically show increased allele frequency in farmed compared to wild Atlantic salmon (Table 1). These SVs disrupt or ablate coding genes with diverse functions, including male fertility (e.g. *CATSPERB*^40^), immunity (e.g. B cell survival and signalling, *GIMAP8*^41^ and two distinct *CD22*^42^ genes), circadian control of metabolism (*NR1D2*^43^), lipid metabolism and insulin sensitivity (*ELOVL6*^44^), and melanin transport and deposition (*MYRAP*^45^) (Table 1). We observed four deletions that disrupt conserved lncRNAs of unknown function, and several large SVs that cover multiple genes, for instance a 423 kb inversion on Chromosome 7 containing 16 genes that was absent in 257 wild salmon (Table 1). In summary, this data demonstrates that diverse gene functions beyond neurological and behavioural pathways were altered by the domestication of Atlantic salmon due to altered selective pressure or drift.

**Table 1.** Major effect SVs under divergent selection in farmed and wild Atlantic salmon.

| Chr | Start | Size | Type | Impact | SV genotype frequencies |  |  |  |  |  |  |
| --- | --- | --- | --- | --- | --- | --- | --- | --- | --- | --- | --- |
|  |  |  |  |  | F<sub>ST</sub> | 0/0<br>Wild | 0/0<br>Farmed | 0/1<br>Wild | 0/1<br>Farmed | 1/1<br>Wild | 1/1<br>Farmed |
| 1 | 15,177,232 | 23,362 | DEL | Deletes coding exons 3-12 in metabolic gene *SCCPDH* (LOC106569909, 12 exons) and lncRNA conserved in teleosts (LOC106569968) | 0.12 | 0.95 | 0.76 | 0.05 | 0.24 | 0.00 | 0.00 |
| 1 | 15,282,772 | 9,209 | DUP | Duplicates coding exons 5-10 within immune gene *GIMAP8* (LOC106569455, 14 exons) | 0.10 | 1.00 | 0.94 | 0.00 | 0.06 | 0.00 | 0.00 |
| 1 | 38,534,900 | 2,471 | DEL | Deletes coding exons 15-16 within sperm motility gene *CATSPERB* (106602505, 26 exons) | 0.11 | 1.00 | 0.91 | 0.00 | 0.09 | 0.00 | 0.00 |
| 1 | 53,229,610 | 801,673 | DUP | Duplicates region containing 9 coding genes, including immune gene *Pentraxin* (LOC100136583) | 0.27 | 0.96 | 0.65 | 0.04 | 0.32 | 0.00 | 0.03 |
| 1 | 63,072,912 | 1,133 | DEL | Deletes coding exons 16-17 within cell fusion gene *ADAM12* (LOC106607406, 23 exons) | 0.15 | 1.00 | 0.94 | 0.00 | 0.03 | 0.00 | 0.03 |
| 1 | 134,577,173 | 742 | DEL | Deletes lncRNA conserved in salmonids (LOC106567697) | 0.28 | 0.95 | 0.68 | 0.05 | 0.26 | 0.00 | 0.06 |
| 2 | 8,188,202 | 8,134 | DEL | Deletes coding exons 5-10 within glycoprotein gene *TUFT1* (LOC106575489, 16 exons) | 0.12 | 0.98 | 0.85 | 0.02 | 0.15 | 0.00 | 0.00 |
| 2 | 15,507,544 | 2,071 | DUP | Duplicates coding exons 12-15 within *HMCN1* (LOC106578676, 19 exons) | 0.24 | 0.28 | 0.00 | 0.16 | 0.00 | 0.56 | 1.00 |
| 2 | 45,905,818 | 49,351 | DEL | Deletes coding exons 1-25 of cellular adhesion gene *ITGAL* (106588084, 29 exons) | 0.11 | 0.95 | 0.76 | 0.05 | 0.24 | 0.00 | 0.00 |
| 2 | 51,645,286 | 1,172 | DEL | Deletion within coding exon 9 (frameshift) of endocytosis gene *SMAP1* (LOC100286439, 10 exons) | 0.15 | 1.00 | 0.91 | 0.00 | 0.09 | 0.00 | 0.00 |
| 3 | 53,262,801 | 56,833 | DUP | Disrupts coding sequence and intergenic region of two tandem *HEBP2* genes (LOC106600932, LOC106600932) | 0.19 | 0.96 | 0.79 | 0.04 | 0.12 | 0.00 | 0.09 |
| 4 | 33,772,841 | 2,115 | DEL | Deletes coding exons 21-26 of *PCNX1* (LOC106602984, 32 exons) | 0.24 | 1.00 | 0.91 | 0.00 | 0.03 | 0.00 | 0.06 |
| 5 | 23,514,943 | 157 | DEL | Deletes coding exon 8 of *PIGG* isoform 2 (LOC106604548, 8 exons) causing a frameshift | 0.35 | 1.00 | 0.76 | 0.00 | 0.24 | 0.00 | 0.00 |
| 5 | 29,459,708 | 1,886 | DEL | Deletes coding exons 2-3 within GTPase-activating gene *TBC1D2* (LOC106604634, 16 exons) | 0.10 | 1.00 | 0.94 | 0.00 | 0.06 | 0.00 | 0.00 |
| 5 | 54,982,436 | 5,313 | DUP | Affecting coding exons 6-8 within circadian regulator gene *NR1D2* (LOC100136378, 8 exons). Introduces stop codon | 0.15 | 0.84 | 0.50 | 0.10 | 0.38 | 0.06 | 0.12 |
| 6 | 1,542,320 | 19,710 | DUP | Duplicates coding exons 5-7 within immune gene *CD22* (106606237/8, 8 exons) | 0.13 | 0.87 | 0.62 | 0.10 | 0.29 | 0.03 | 0.09 |
| 6 | 29,579,766 | 5,320 | DEL | Deletes lncRNA conserved in salmonids (LOC106607070) | 0.20 | 0.85 | 0.53 | 0.14 | 0.35 | 0.01 | 0.12 |
| 7 | 21,191,252 | 422,735 | INV | Inverts region containing 16 coding genes | 0.11 | 1.00 | 0.91 | 0.00 | 0.09 | 0.00 | 0.00 |
| 9 | 21,282,095 | 11,299 | DUP | Duplicates coding exon 2 within *PGBD3* (LOC106611080, 4 exons) | 0.12 | 0.99 | 0.91 | 0.01 | 0.06 | 0.00 | 0.03 |
| 9 | 53,275,027 | 100,799 | DUP | Fusion of region containing last 10 coding exons of *TAPT1* (LOC106611550) with first 4 coding exons of *PROM1* (LOC106611549) | 0.15 | 0.84 | 0.56 | 0.12 | 0.29 | 0.03 | 0.15 |
| 10 | 23,225,394 | 32,774 | DEL | Deletes region containing six tRNA genes | 0.14 | 0.99 | 0.85 | 0.01 | 0.15 | 0.00 | 0.00 |
| 11 | 13,465,612 | 5,950 | DEL | Deletes exon 1 within lncRNA conserved in teleosts (LOC106562070, 3 exons) | 0.10 | 1.00 | 0.94 | 0.00 | 0.06 | 0.00 | 0.00 |
| 12 | 21,083,103 | 1,693 | DEL | Deletes coding exon 2-3 within uncharacterised gene (LOC106564648, 6 exons) | 0.25 | 0.96 | 0.71 | 0.04 | 0.24 | 0.00 | 0.06 |
| 14 | 14,287,987 | 18,976 | DUP | Duplicates coding exons 8-15 within melanosome transport gene *MYRIP* (LOC106568916, 15 exons) | 0.36 | 0.96 | 0.62 | 0.02 | 0.24 | 0.02 | 0.15 |
| 14 | 83,617,466 | 91,512 | DUP | Duplicates region containing 9 coding exons from *FAM126A* (LOC106570580), complete cytokine gene *IL6* (LOC106570581) and coding exon 1 from *RAPGEF5* (LOC106570584) | 0.13 | 0.98 | 0.88 | 0.02 | 0.06 | 0.00 | 0.06 |
| 18 | 56,889,482 | 39,099 | DUP | Duplicates coding exons 1-12 within immune gene *CD22* (LOC106577812, 20 exons) | 0.12 | 0.94 | 0.76 | 0.05 | 0.18 | 0.01 | 0.06 |
| 18 | 64,338,324 | 852 | DEL | Deletes coding exon 7 within gene *PARP14*-like (LOC106578007, 7 exons) and ablates stop codon | 0.15 | 0.84 | 0.56 | 0.14 | 0.32 | 0.02 | 0.12 |
| 19 | 51,422,161 | 31,121 | INV | Flips coding exon 1-2 within fatty acid elongation gene *ELOVL6* (LOC106579283, 4 exons) | 0.11 | 0.93 | 0.71 | 0.07 | 0.29 | 0.00 | 0.00 |
| 22 | 40,200,901 | 5,863 | DEL | Deletes coding exon 2 within *PLEKHA6* (LOC106583501, 24 exons) | 0.13 | 0.97 | 0.85 | 0.02 | 0.06 | 0.01 | 0.09 |
| 24 | 11,833,364 | 165 | DUP | Deletes half of coding exon 2 within tRNA methyltransferase gene *TRMT2A* (LOC106584929, 12 exons) | 0.11 | 0.09 | 0.32 | 0.44 | 0.47 | 0.46 | 0.21 |
| 24 | 19,661,320 | 266,147 | INV | Affects 6 coding genes, inverting 5 genes completely and all but first exon of *AAK1* (LOC106585601) | 0.16 | 0.83 | 0.44 | 0.17 | 0.56 | 0.00 | 0.00 |
| 27 | 42,220,948 | 341 | DEL | Partially deletes exon 4 in angiogenesis gene (*ANG2* LOC106589146, 5 exons) causing frameshift | 0.12 | 0.56 | 0.24 | 0.34 | 0.50 | 0.10 | 0.26 |
| 28 | 3,887,040 | 5,373 | DUP | Duplication affecting zinc transporter gene *SLC39A11* (LOC100380452, 10 exons) causing frameshift | 0.14 | 0.95 | 0.79 | 0.04 | 0.09 | 0.02 | 0.12 |
| 28 | 16,046,880 | 24,780 | DUP | Fusion involving coding exons 9-16 of sodium transport gene *SLC38A10* (LOC106589592, 16 exons) and exons 1-3 of vesicular transport gene *TEPSIN* (15 exons) | 0.52 | 0.86 | 0.29 | 0.10 | 0.26 | 0.04 | 0.44 |

Genotypes: 0/0: homozygous lacking SV; 0/1 heterozygous for SV 1/1 homozygous for SV

## Discussion

Despite an increasing shift towards the use of long-read sequencing for SV discovery^1,2^, these technologies remain prohibitively expensive for large-scale population genetics, making such datasets scarce in most species. Consequently, it remains a timely challenge to extract reliable SV calls from the more extensive repository of short-read genome sequencing datasets, which continue to emerge rapidly in many species, largely for use in SNP analyses. The approach reported can be applied for reliable SV detection and genotyping using such data in any species with a reference genome. A critical step - unique to this study - was the curation of all SV calls using SV-plaudit^10^. This approach demands significant manual effort, equivalent to approximately two weeks for a small team of trained curators, yet was efficient in retaining predominantly true calls, and allowed us to demonstrate the value of filtering complex regions to drastically reduce the FDR. The overall extreme FDR for SV discovery advocates for the routine application of such curation in SV studies based on short-read sequencing, particularly if ‘gold-standard’ SVs defined by past work are unavailable.

The SVs reported provide a novel resource for future studies on the genetic architecture of traits in Atlantic salmon, which has excluded SVs until now. It will be useful to overlap our SVs with genomic regions of interest such as QTLs defined by SNPs, to investigate SVs as putative causal variants. For example, we discovered a duplication on chromosome 14 that likely destroys the function of the *MYRIP* gene, which is involved in melanosome transport^45^ – a past study discovered a single QTL on chromosome 14 that explained differences in melanocyte pigmentation between wild and domesticated fish^46^, which may be linked to this newly discovered SV. It will also be useful in future studies to apply SV markers directly in genome wide association analyses, and to test their value for genomic prediction in salmon breeding programmes^11,12^. While our study captured hundreds of Atlantic salmon genomes representing several major phylogeographic groups, it fails to capture broader genetic diversity within this species, and due to the retention of only high confidence SV calls, our method may be prone to false negatives. Further, inherent limitations of short read sequencing data for SV detection presumably obscures detection of many SVs, suggesting future SV studies in Atlantic salmon must also focus on adapting long-read sequence data, and integrating short and long-read data for optimal SV discovery^1^.

We discovered intact pTSsa2 polymorphisms within our SV dataset, and provided evidence for transposon expansion after the split of *S. salar* and *trutta* ∼10 Mya^16^ (Fig. 2). The pTSsa2 transposon appears with high copy number in the Atlantic salmon genome, suggesting an important role in shaping very recent genome architecture. Transposons have largely been excluded from studies of contemporary genetic variation in salmonids, but were central to genome rediploidization after the Ss4R WGD^17^, and likely contributed to the evolution of the sex determining locus, e.g.^47^. As work in other taxa has revealed that transposon polymorphisms contribute to adaptive evolution ^48,49^ and speciation^50^, future studies on pTSsa2 should investigate such possibilities in *Salmo*. We also showed that Atlantic salmon deletion SVs are more likely to overlap genes retained as ohnolog pairs from the Ss4R WGD event compared to singleton genes. This supports the hypothesis that WGD events buffer against potential deleterious impacts of SVs on gene function and regulation, consistent with past work^29,51^. However, the link between SVs and the Ss4R WGD requires further investigation to more fully dissect the role of selection and drift in driving SV retention.

We discovered many SVs showing genetic divergence between farmed and wild Atlantic salmon linked to synaptic genes responsible for behavioural variation^38,39^. Most were rare alleles in wild fish and showed a small to moderate increase in frequency in domesticated populations, consistent with a polygenic genetic architecture for behavioural traits altered by domestication, including risk-taking behaviour, aggression, and boldness^32,52,53,54,55,56^, affecting many unique genes from the same functional networks, mirroring the polygenic basis for many human neurological traits^57,58,59^. The disruption of mammalian orthologs for many of the same synaptic genes cause disorders including schizophrenia, intellectual disability, autism, and Alzheimer’s (Supplementary Table 10). For Atlantic salmon, we did not establish if these SVs are causative variants or in linkage disequilibrium with other variants under selection. In several cases, it is likely that the SVs discovered are causative variants due to their disruptive nature on protein coding gene sequence potential (e.g. Table 1), including the ablation of the key synaptic protein neurexin-2, which caused autism-related behaviours when induced experimentally in mice^60^. However, as many of the outlier SVs were located in non-coding regions, this points to regulatory effects on gene expression, which may have minor or additive effects on behavioural traits. Future work should test whether the outlier SVs alter the expression or function of synaptic genes and directly influence behavioural phenotypes. Beyond neurological systems, domestication altered the frequencies of numerous major effect SVs disrupting genes with diverse functional roles (Table 1), providing candidate causative variants for ongoing investigations into diverse traits. For instance, an increased frequency of SVs ablating the *ELOVL6* and *NR1D2* genes in domesticated fish, which play key roles in lipid metabolism, insulin resistance, and the coordination of metabolic functions with the circadian clock^44,45^, is highly consistent with a recent transcriptomic study demonstrating altered metabolism linked to disrupted circadian regulation in domesticated compared to wild Atlantic salmon^61^.

To conclude, given the rapidly growing recognition of the importance of establishing the role of SVs in adaptation and other evolutionary processes in natural populations^62,63^, in addition to commercial variation relevant to breeding of farmed animals^64,65^, we anticipate that this reliable description of the SV landscape in Atlantic salmon will encourage more studies exploiting SV markers to address both fundamental and applied questions in the genetics of non-model species.

## Methods

### Sequencing data

Paired-end whole genome sequencing data (mean 8.1x coverage, 2 x 100-150 bp) was generated for 472 Atlantic salmon on several different platforms (Supplementary Table 1). DNA extraction, quality control and sequencing library preparation followed standard methods. Wild Atlantic salmon were sampled either during organized fishing expeditions or by anglers during the sport fishing season with DNA extracted from scales. We sampled n=80 wild Canadian individuals from 8 sites, n=359 Norwegian individuals from 52 sites (including n=5 landlocked dwarf salmon), n=8 Baltic individuals from a single site and n=4 White sea individuals from a single site. Whole genome sequencing data was generated for 21 farmed individuals (n=12 individuals from Mowi ASA; n=9 samples from Xelect Ltd) and downloaded for a further 20 individuals (NCBI accession: PRJNA287458).

### SV detection and genotyping

Sequence alignment to the unmasked ICSASG_V2 assembly (GCA_000233375.4)^17^ was done using BWA v0.7.13^66^. Reads were mapped to the complete reference, including unplaced scaffolds, with random placement of multi-mapping reads^67^. Reads mapping to unplaced scaffolds were discarded. Alignments were converted to BAM format in Samtools v0.1.19^68^. Alignment quality, batch effects and sample error were further assessed using Indexcov goleft v0.2.1^69^. Gap regions were extracted and converted to BED format using a Python script (Supplementary Note 1); SV calls overlapping these regions were identified using Bedtools^70^ and removed. Sample coverage was estimated using mosdepth v0.2.3^71^. High depth regions were defined as ≥100x coverage and removed; this cut-off was a compromise to avoid generating too many false SV calls, balanced against the risk of losing real SVs. High depth regions located within 100bp were merged. SV detection was done using the Lumpy-based tool Smoove V.2.3^21^ with genotypes called by SVtyper^22^. Gap and high-depth regions were combined into a single BED file, which can optionally be used to exclude these locations from SV detection in Lumpy (-exclude option). All of the above steps were combined in a Snakemake (v.3.11.0)^20^ workflow, with the input being paired-end sequencing data (FASTQ format), and the output a VCF file with SV locations and genotypes for all individuals in a study (Supplementary Figure 1; Supplementary Note 2 provides Snakefile).

### SV-plaudit curation

All 165,116 SV calls generated in the study were curated using SV-plaudit^10^. A plotCritic website was setup on Amazon Web Services where variant images produced in samplot v1.01 (https://github.com/ryanlayer/samplot) were deployed. SV curation involved the random visualization of one homozygous wild-type (0/0; lacking SV, identical to reference genome), two heterozygous (0/1, with one SV copy) and two homozygous-alternate (1/1, with two SV copies) individuals per SV, done using cyvcf2 v0.11.5^72^. With each image the question “is this variant real?” was answered (options: ‘No’, ‘Yes’, or ‘Maybe’). Only high confidence variants (‘Yes’) were kept for downstream analysis. Three different co-authors (ACB, MKG, EP) team-curated the full SV set. 1,000 random plots were commonly curated by each researcher to establish congruence in decision making, and there was 100% agreement concerning high confidence (‘Yes’) variants. Subsequently the SV plots were divided randomly and each set validated independently across the 3 researchers and then merged.

### SV annotation

High confidence SVs retained following SV-plaudit curation were filtered to remove redundant SVs using the Bedtools *intersect* function (90% reciprocal overlap), removing 133 SVs and leaving 15,483 SVs used in further analysis (Supplementary Data 2). The association between SVs and RefSeq genes within the ICSASG_v2 assembly was done using SnpEff^26^ (default parameters). GO enrichment tests were done using the ‘weight01’ algorithm and Fisher’s test statistic in the TopGo package^73^. The background set was all genes in the RefSeq annotation. The R package ‘Ssa.RefSeq.db’ (https://gitlab.com/cigene/R/Ssa.RefSeq.db)^74^ was used to retrieve GO annotations from the ICSASG_v2 genome. The overlap between SV locations and repeats in the ICSASG_v2 annotation was done using Bedtools^61^ against an existing database^17^.

### Phylogenetic analyses

pTSsa2 sequences including EF685967 were used in BLASTn^75^ searches against the NCBI nucleotide database (restricted to Salmonidae) in addition to unmasked assemblies for Atlantic salmon (ISCASG_v2), brown trout (GCA_901001165.1), Arctic charr (GCA_002910315.2), rainbow trout (GCA_002163495.1), chinook salmon (GCA_002872995.1) and coho salmon (GCA_002021735.2). Sequence alignments were performed using Mafft^76^ with default settings. Phylogenetic analysis was done using the IQTREE server^77^ with estimation of the best-fitting nucleotide substitution model (Bayesian Information Criterion) and 1,000 ultrafast bootstraps^78^.

### SV validation by MinION sequencing

PCR primers are shown in Supplementary Table 3. PCRs were performed using LongAmp® Taq (New England Biolabs) with 1 cycle of 94°C for 30s, 30 cycles of 94°C for 30s, 56°C for 60s and 65 °C for 50s/kb, followed by a 10 min extension at 65°C. Amplicons for different SVs in each fish individual were pooled and cleaned using AMPure XP beads (Beckman Coulter). 250ng pooled DNA was used to create sequencing libraries with a 1D SQK-LSK109 kit (Oxford Nanopore Technologies, ONT). DNA was end-repaired using the NEBNext Ultra II End Repair/dA Tailing kit (New England Biolabs) and purified using AMPure XP beads. Native barcodes were ligated to end-repaired DNA using Blunt/TA Ligation Master Mix. Barcoded DNA was purified with AMPure XP beads and pooled in equimolar concentration to a total of 200 ng per library (∼0.2 pmol). AMII Adapter mix (ONT) was ligated to the DNA using Blunt/TA Ligation Master Mix (New England Biolabs) before the adapter-ligated library was purified with AMPure XP beads. DNA concentration was determined at each step using a Qubit fluorimeter (Thermo Fisher Scientific) with a ds-DNA HS kit (Invitrogen).

Sequencing libraries were loaded onto MinION FLO-MIN106D R9.4.1 flow cells (ONT) and run via MinKNOW for 36h without real-time basecalling. Basecalling and demultiplexing was performed with Guppy v2.3.7. FASTQ files were uploaded into Geneious Prime 2019.1.1 and simultaneously mapped to a reference of sequences spanning all candidate SV regions in the ISCSAG_v2 assembly. Mapping was done with the following parameters: ‘medium-fast sensitivity’, ‘finding structural variants’, including ‘short insertions’ and ‘deletions’ of any size, with the setting ‘map multiple best matches’ set to ‘None’, and the minimum support for SV discovery set to 2 reads. Alignments were inspected for the presence and genotype of the SV. Amplicons with <50x coverage to the target SV region were discarded as failed PCRs. When alignments matched the predicted SV breakpoints and size, the SV call was considered correct. When >90% of the aligned reads matched to the expected SV and breakpoints (i.e. a gap for deletions, an insertion for duplications and flipped reads for inversions compared to the reference) it was classified 1/1 homozygous. When at least 10% of the aligned reads matched to both the reference genome state, in addition to the 1/1 state, the locus was classified 0/1 heterozygous.

### Association between SVs and Ss4R ohnologs

The code used to identify a genome-wide set of Ss4R ohnologs, along with a description of the genome assembly annotations employed, is available at https://gitlab.com/sandve-lab/salmonid_synteny and https://gitlab.com/sandve-lab/defining_duplicates. Orthogroups were constructed with Orthofinder^79^ using seven salmonid species (Atlantic salmon, rainbow trout, Arctic charr, coho salmon, huchen *Hucho hucho*, and European grayling *Thymallus thymallus*), five additional actinopterygians (zebrafish, medaka *Oryzias latipes*, northern pike *Esox lucius*, three-spined stickleback *Gasterosteus aculeatus* and spotted gar *Lepisosteus oculatus*), and two mammals (human and mouse *Mus musculus*). For each orthogroup, we extracted nucleotide protein coding sequences, aligned them with Macse^80^ and built gene trees using TreeBeST^81^. Trees were split into smaller subtrees at the node representing the divergence between pike and salmonids. To derive a final set of Atlantic salmon Ss4R ohnologs, we used both synteny and gene tree topology criteria. Firstly, we required that the subtrees branched with northern pike as the sister to salmonids and outgroup to Ss4R^16,17^ and contained either exactly two (ohnologs) or exactly one (singletons) Atlantic salmon genes. Secondly, we removed any putative Ss4R ohnologs falling outside conserved synteny blocks predicted using iadhore^82^. A final set of ohnolog pairs is provided in Supplementary Data 4, which contains all gene trees in NWK format.

We used the *fisher.exact()* function in R to compare the observed counts of SVs overlapping singleton and ohnologs with the total counts of singletons and ohnologs. To test for association between ohnolog expression divergence and SV overlap, we used a 15 tissue RNA-Seq dataset^17^ available as a TPM (transcripts per million reads) table in the salmofisher R-package https://gitlab.com/sandve-lab/salmonfisher. We used the *cor()* function in R to compute median Spearman’s tissue expression correlation for all ohnolog pairs where one copy was overlapped by an SV. We then computed median correlations for 1,000 randomly sampled ohnolog sets of the same size. The *P*-value was estimated as the proportion of resampled medians lower than the observed median for ohnologs overlapped by SVs. Tests comparing expression level between genes that were either overlapped or not overlapped by SVs were conducted using the sum log10 transformed TPM for each gene across all 15 tissues. The function *wilcox_test* within the R-package *rstatix* was used to calculate *P*-values for differences in expression levels. The code used is available at https://gitlab.com/ssandve/atlantic_salmon_sv_ohnolog_analyses/.

### Association of SVs with brain ATAC peaks

Four Atlantic salmon (freshwater stage, 26-28g) were killed using a Schedule 1 method following the Animals (Scientific Procedures) Act 1986. Around 50mg homogenized brain tissue was processed to extract nuclei using the Omni-ATAC protocol for frozen tissues^83^. Nuclei were counted on an automated cell counter (TC20 BioRad, range 4-6 um) and further confirmed intact under microscope. 50,000 nuclei were used in the transposition reaction including 2.5 µL Tn5 enzyme (Illumina Nextera DNA Flex Library Prep Kit), incubated for 30 minutes at 37 °C in a shaker at 200 rpm. The samples were purified with the MinElute PCR purification kit (Qiagen) and eluted in 12μL elution buffer. qPCR was used to determine the optimal number of PCR cycles for library preparation^84^ (8-10 cycles used). Sequencing libraries were prepared with short fragments and fragments >1,000 bp removed using AMPure XP beads (Beckman Coulter, Inc.). Fragment length distributions and confirmation of nucleosome banding patterns were determined on a 2100 Bioanalyzer (Agilent) and the library concentration estimated using a Qubit system (Thermo Scientific). Libraries were sent to the Norwegian Sequencing Centre, where paired-end 2 x 75 bp sequencing was done on an Illumina HiSeq 4000. The raw sequencing data is available through ArrayExpress (Accession: E-MTAB-9001).

ATAC-Seq reads were aligned to the Atlantic salmon genome (ICSASG_v2) using BWA (v0.7.17)^66^ and a merged peak set called combining the four replicates using Genrich (https://github.com/jsh58/Genrich) with default parameters, apart from “-m 20 -j” (minimum mapping quality 20; ATAC-Seq mode). Bedtools was used to identify SVs overlapping ATAC-Seq peaks (filtered at corrected *P* ≤0.01) associated to genes, defined as being located within 3,000 bp up/downstream of the start and end coordinates of the longest transcript per gene.

### Population structure analyses and F_ST_ analyses

PCAs were performed separately on the complete set of high confidence deletions (14,017), duplications (1,244) and inversions (242) using the *prcomp* and *autoplot* functions within GGplot2^85^ in R. Genotypes were coded into bi-allelic marker format to be compatible with standard population genetics methods. Population structure was examined using NGSadmix^86^ tested for group sizes of K=2-4.

F_ST_ values were calculated for all high confidence SVs using VCFtools v0.1.16^87^ with the Weir and Cockerham method^37^ comparing 34 Norwegian farmed vs. 257 Norwegian wild Atlantic salmon (Fig. 4a provides rationale for sample selection). To establish the significance of each F_ST_ value, individuals from the two groups were randomly split into two sets of the original size (i.e. 34 vs. 257 individuals) 200 times, before the distribution of resultant F_ST_ values was plotted using the ggplot2 function *geom_freqpoly* (binwidth = 0.01). Per SV *P*-values were considered as the proportion of F_ST_ values obtained in the 200 random distributions higher than the F_ST_ in the observed distribution. Thus, if 10/200 randomly sampled F_ST_ values above the observed F_ST_ value were recorded, *P*=0.05 was assigned. We further applied an F_ST_ cutoff to include SVs where 99.7% of all F_ST_ values fell above the randomly sampled values (F_ST_ >0.103). Any SVs lacking alternative alleles in the compared groups were excluded. Code to perform these analyses is provided in Supplementary Note 3.

### Annotation of SV outliers

GO enrichment tests for genes linked to the SV outliers (*P <*0.05) were done as described in the section ‘SV annotation’, with the background gene set restricted to all RefSeq genes linked to SVs by SnpEff. To investigate the expression of genes linked to SV outliers, we used existing RNA-seq data^17^, representing normalized counts per million (CPM) for 10 tissues (brain, liver, muscle, spleen, pancreas, heart, pyloric, gill, skin and foregut). We filtered any genes where the across-tissue sum of CPM was <1.0. A ‘tissue specificity’ index was calculated, representing the sum across-tissue CPM divided by the CPM per tissue. We tested whether genes linked to SV outliers by SnpEff, in addition to a subset contributing to significant GO terms (*P*<0.01), differed from the transcriptome-wide expectations. Hypergeometric tests were used (*dhyper* function in R) to compare the number of genes in the two gene sets with a tissue specificity index ≥0.5 compared to all genes in the transcriptome. Two-sample t-tests (*t.test* function in R) were used to compare differences in mean CPM between the two gene sets compared to all genes in the transcriptome. BLAST was used to cross-reference protein products of genes linked to SV outliers against 3,840 unique proteins detected in the zebrafish synaptic proteome^38^ (downloaded from the GRCz11 assembly version using BioMart at Ensembl.org), taking forward the top zebrafish BLAST hit (cut-off: 40% identity, 40% query coverage) as a query in a reciprocal BLAST against all *S. salar* RefSeq proteins (no cut-off); evidence for orthology was accepted when the candidate zebrafish protein showed a best hit to the original query in the complete salmon proteome. We used the *fisher.exact()* function in R to test if the 584 significant F_ST_ outlier SVs were more likely to overlap brain ATAC-Seq peaks than non-significant SVs (*P* > 0.05), which was done considering all expressed genes (TPM ≥1) in the RNA-Seq tissue atlas described above^17^ and a subset of the same genes most highly expressed in brain (filtered for genes where brain was among the top 3 tissues for TPM). The bedtools^61^ intersect function was used to associate ATAC-Seq peaks with SVs. The code used is available at https://gitlab.com/ssandve/atlantic_salmon_sv_ohnolog_analyses/.

### Data availability

New genome sequences generated are available through the European Nucleotide Archive (project accession: PRJEB38061, released upon publication). Sample accession numbers for all 492 Atlantic salmon genomes are provided in Supplementary Table 1 (available upon publication). ATAC-Seq reads were deposited in ArrayExpress (accession: E-MTAB-9001).

### Code availability

Python script used to identify regions in ICSASG_v2 genome and convert output to BED file: Supplementary Note 1.

Snakefile and associated code for SV detection pipeline: Supplementary Note 2.

R script used to obtain F_ST_ values from random comparisons and establish probability value for outlier SVs: Supplementary Note 3.

Code to define orthogroups and build gene trees: https://gitlab.com/sandve-lab/salmonid_synteny.

Code to identify Atlantic salmon ohnolog pairs from ortholog groups and gene trees: https://gitlab.com/sandve-lab/defining_duplicates.

Code to analyse overlaps between SVs, ohnologs and ATAC-Seq data: https://gitlab.com/ssandve/atlantic_salmon_sv_ohnolog_analyses/.

## Acknowledgements

The study was supported by Biotechnology and Biological Sciences Research Council grants BB/M016455/1, BB/S004181/1, BBS/E/D/10002070 and BBS/E/D/30002275. Wild Atlantic salmon genome sequencing was funded by the Research Council of Norway (The Aqua Genome project; ref: 221734). We thank Dr Chris Hollenbeck (formerly Xelect Ltd; currently Texas A&M, USA) for support with Snakemake and Drs Serap Gonen and Matt Baranski (Mowi AS) for sharing sequencing data. We thank Terese Andersstuen and Dr Mariann Árnyasi (both NMBU) for organizing the sequencing of wild Atlantic salmon samples. We acknowledge the use of computing clusters at the University of Aberdeen (Maxwell), University of Edinburgh (EDDIE) and CIGENE, NMBU (Orion). Storage resources were provided by the Norwegian National Infrastructure for Research Data (NIRD, project NS9055K). DJM thanks Prof. Seth Grant (University of Edinburgh) for helpful discussion concerning the Atlantic salmon SV outliers and synaptic genes.

## Contributions

DJM, SL, IAJ and SRS conceived the study. ACB, RL and TN developed the SV detection workflow. ACB performed downstream analyses with contributions from MKG, DR, DJM, TDM, SRS, and EP. DJM (lead supervisor), TJA, IAJ and SAM supervised ACB. ACB and MDG performed MinION sequencing. SLL and MHH performed ATAC-Seq. KH, HS., BF-L, JE, CRP and LB provided wild Atlantic salmon samples. ACB, DJM and SRS drafted the text and figures. All authors commented on and approved the final manuscript.

## Supplementary information

Supplementary Figure 1. Snakemake pipeline for end-to-end SV detection.

Supplementary Figure 2. Locations of complex regions in Atlantic salmon genome.

Supplementary Figure 3. Example of SV-plaudit image for a high confidence deletion SV call

Supplementary Figure 4. Example of SV-plaudit image for a false positive deletion SV call excluded from further analyses.

Supplementary Figure 5. Example of SV-plaudit image for a high confidence duplication SV call retained in further analyses

Supplementary Figure 6. Example of SV-plaudit image for a false positive duplication SV call excluded from further analyses

Supplementary Figure 7. Example of SV-plaudit image for a high confidence inversion SV call retained in further analyses

Supplementary Figure 8. Example of SV-plaudit image for a false positive inversion SV call excluded from further analyses

Supplementary Figure 9. SV sizes before SV-plaudit curation

Supplementary Figure 10. Sequencing depth was not a strong predictor of the final number of high-confidence SVs retained after SV-plaudit curation.

Supplementary Figure 11. Summary of 168 SV regions used for MinION amplicon sequencing to validate Lumpy/SVtyper SV and genotype calls.

Supplementary Figure 12. Example of congruence between SV/genotype calls and data generated by MinION amplicon sequencing.

Supplementary Figure 13. Example of congruence between SV/genotype calls and data generated by MinION amplicon sequencing.

Supplementary Figure 14. Example of congruence between SV/genotype calls and data generated by MinION amplicon sequencing.

Supplementary Figure 15. PCAs showing the same data presented in Fig. 1g-i (main text), except visualized according to latitude

Supplementary Figure 16. PCA analyses done on SV genotype calls prior to SV-plaudit curation

Supplementary Figure 17. SV annotation by SnpEff

Supplementary Figure 18. Overlap between high confidence SVs and protein coding genes in the ICSASG_v2 annotation.

Supplementary Figure 19. Number of high impact annotations per snpEff effect for high-confidence Atlantic salmon SVs.

Supplementary Figure 20. Maximum likelihood tree presented in Fig. 2 including sample identifiers, genomic locations of pTSsa2 sequences and bootstrap values.

Supplementary Figure 21. Circos plot showing the genomic locations of pTSsa2 sequences in the Atlantic salmon genome

Supplementary Figure 22. Expression characteristics of ohnologs depending on SV overlap. Supplementary Figure 23. Tissue expression levels comparing SV outliers with transcriptome wide expectations for nine tissues

Supplementary Figure 24. Tissue specificity comparing SV outliers with transcriptome wide expectations for nine tissues

Supplementary Figure 25. Heatmap showing individual SV genotypes for 45 SV outliers linked to synapse genes.

Supplementary Note 1. Python script used to extract gap regions in the ICSASG_v2 genome and and convert the outputs to a BED file

Supplementary Note 2. Snakefiles and associated code for SV calling pipeline

Supplementary Note 3: Custom R script used to obtain F_ST_ values from random comparisons and establish probability value for outlier SVs

Supplementary Table 1. Details of samples used in study

Supplementary Table 2. SV call statistics per individual across 492 Atlantic salmon samples following different filtering steps

Supplementary Table 3. Validation of SV calls and genotypes using MinION sequencing

Supplementary Table 4. GO Biological Process enrichment analysis for genes affected by high impact deletions, duplications and inversions

Supplementary Table 5. Genes contributing to significant GO terms for high impact SVs

Supplementary Table 6. Fishers Exact test results contrasting the overlap between SVs with singleton genes vs. Ss4R ohnolog genes.

Supplementary Table 7. GO enrichment analysis for genes linked to SV outliers between wild and farmed Atlantic salmon

Supplementary Table 8. Genes contributing to significant GO terms for genes linked to SV outliers.

Supplementary Table 9. Statistical tests of two expression characteristics (specificity and level) across a panel of tissues for 327 SV outlier linked genes contributing to significantly enriched GO biological processes in comparison to a transcriptome-wide set gene set.

Supplementary Table 10. Detailed annotation of prioritized SV outliers between farmed and wild Atlantic salmon linked to genes with synaptic functions.

Supplementary Data 1: Full SV dataset and genotypes prior to SV-plaudit curation

Supplementary Data 2: High-confidence SVs retained after SV-plaudit curation, including individual genotypes and SnpEff annotation

Supplementary Data 3: Alignment of SV deletions representing pTSsa2 piggyBac-like DNA transposons (used to Generate Fig. 2)

Supplementary Data 4: High confidence annotation of Ss4R ohnolog and singletons in the Atlantic salmon genome

Supplementary Data 5: Manually filtered SV deletions that alter protein-coding exons

Supplementary Data 6: Significant SV outliers between wild and farmed salmon from Norway

