## Supplementary Information for "The structural variation landscape in 492 Atlantic salmon genomes"

Contents*:*

Supplementary Figure 1. Snakemake pipeline for end-to-end SV detection.
Supplementary Figure 2. Locations of complex regions in Atlantic salmon genome.
Supplementary Figure 3. Example of SV-plaudit image for a high confidence deletion SV call
Supplementary Figure 4. Example of SV-plaudit image for a false positive deletion SV call excluded from further analyses.
Supplementary Figure 5. Example of SV-plaudit image for a high confidence duplication SV call retained in further analyses
Supplementary Figure 6. Example of SV-plaudit image for a false positive duplication SV call excluded from further analyses
Supplementary Figure 7. Example of SV-plaudit image for a high confidence inversion SV call retained in further analyses
Supplementary Figure 8. Example of SV-plaudit image for a false positive inversion SV call excluded from further analyses
Supplementary Figure 9. SV sizes before SV-plaudit curation
Supplementary Figure 10. Sequencing depth was not a strong predictor of the final number of high-confidence SVs retained after SV-plaudit curation.
Supplementary Figure 11. Summary of 168 SV regions used for MinION amplicon sequencing to validate Lumpy/SVtyper SV and genotype calls.
Supplementary Figure 12. Example of congruence between SV/genotype calls and data generated by MinION amplicon sequencing.
Supplementary Figure 13. Example of congruence between SV/genotype calls and data generated by MinION amplicon sequencing.
Supplementary Figure 14. Example of congruence between SV/genotype calls and data generated by MinION amplicon sequencing.
Supplementary Figure 15. PCAs showing the same data presented in Fig. 1g-i (main text), except visualized according to latitude
Supplementary Figure 16. PCA analyses done on SV genotype calls prior to SV-plaudit curation
Supplementary Figure 17. SV annotation by SnpEff
Supplementary Figure 18. Overlap between high confidence SVs and protein coding genes in the ICSASG_v2 annotation.
Supplementary Figure 19. Number of high impact annotations per snpEff effect for high-confidence Atlantic salmon SVs.
Supplementary Figure 20. Maximum likelihood tree presented in Fig. 2 including sample identifiers, genomic locations of pTSsa2 sequences and bootstrap values.
Supplementary Figure 21. Circos plot showing the genomic locations of pTSsa2 sequences in the Atlantic salmon genome
Supplementary Figure 22. Expression characteristics of ohnologs depending on SV overlap.
Supplementary Figure 23. Tissue expression levels comparing SV outliers with transcriptome wide expectations for nine tissues
Supplementary Figure 24. Tissue specificity comparing SV outliers with transcriptome wide expectations for nine tissues
Supplementary Figure 25. Heatmap showing individual SV genotypes for 45 SV outliers linked to synapse genes.

Supplementary Notes

Supplementary Note 1. Python script used to extract gap regions in the ICSASG_v2 genome and and convert the outputs to a BED file

Supplementary Note 2. Snakefiles and associated code for SV calling pipeline
Supplementary Note 3: Custom R script used to obtain Fst values from random comparisons and establish probability value for outlier SVs

*Supplementary Tables attached separately:*

Supplementary Table 1. Details of samples used in study
Supplementary Table 2. SV call statistics per individual across 492 Atlantic salmon samples following different filtering steps
Supplementary Table 3. Validation of SV calls and genotypes using MinION sequencing
Supplementary Table 4. GO Biological Process enrichment analysis for genes affected by high impact deletions, duplications and inversions
Supplementary Table 5. Genes contributing to significant GO terms for high impact SVs
Supplementary Table 6. Fishers Exact test results contrasting the overlap between SVs with singleton genes vs. Ss4R ohnolog genes.
Supplementary Table 7. GO enrichment analysis for genes linked to SV outliers between wild and farmed Atlantic salmon
Supplementary Table 8. Genes contributing to significant GO terms for genes linked to SV outliers.
Supplementary Table 9. Statistical tests of two expression characteristics (specificity and level) across a panel of tissues for 327 SV outlier linked genes contributing to significantly enriched GO biological processes in comparison to a transcriptome-wide set gene set.
Supplementary Table 10. Detailed annotation of prioritized SV outliers between farmed and wild Atlantic salmon linked to genes with synaptic functions.

*Supplementary Datasets attached separately:*

Supplementary Data 1: Full SV dataset and genotypes prior to SV-plaudit curation
Supplementary Data 2: High-confidence SVs retained after SV-plaudit curation, including individual genotypes and SnpEff annotation
Supplementary Data 3: Alignment of SV deletions representing pTSsa2 piggyBac-like DNA transposons (used to Generate Fig. 2)
Supplementary Data 4: High confidence annotation of Ss4R ohnolog and singletons in the Atlantic salmon genome
Supplementary Data 5: Manually filtered SV deletions that alter protein-coding exons
Supplementary Data 6: Significant SV outliers between wild and farmed salmon from Norway

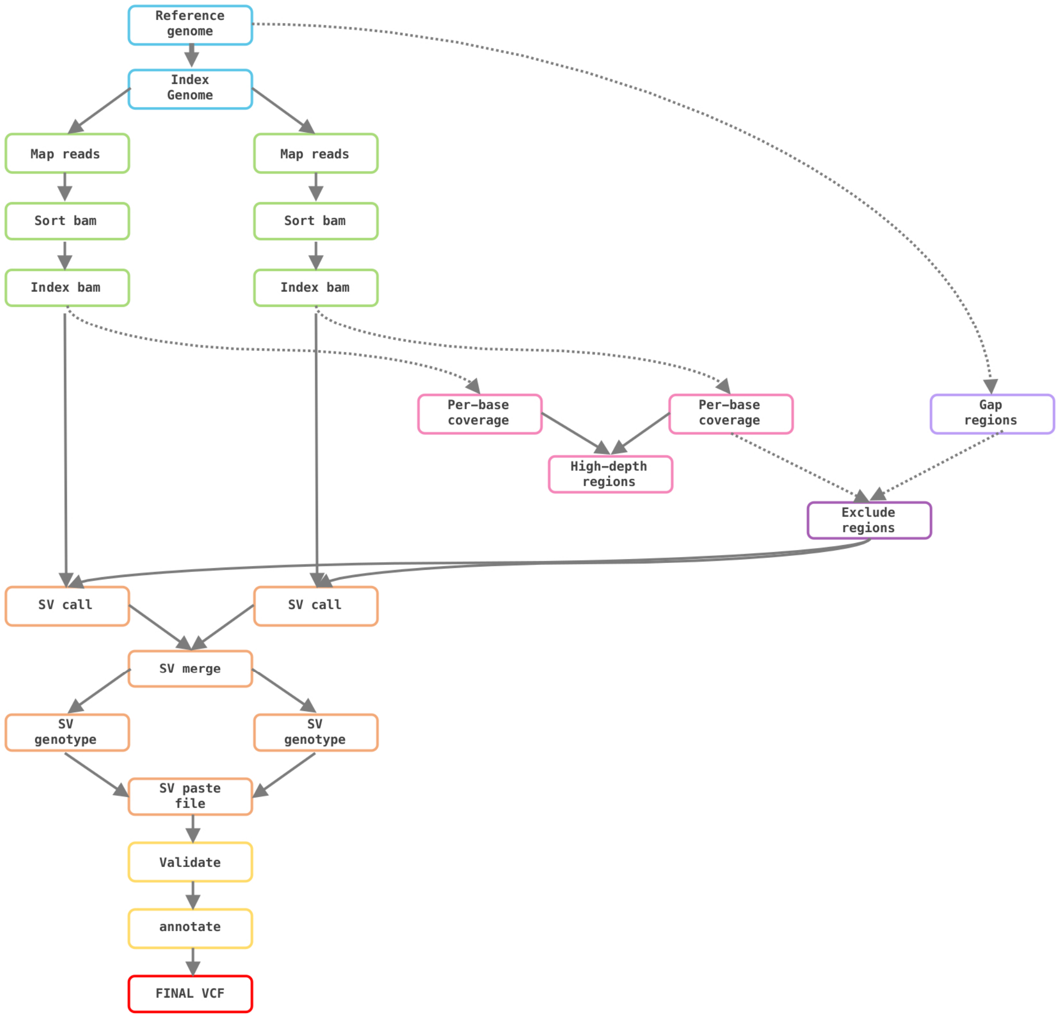


**Supplementary Figure 1.** Snakemake (v.3.11.0) pipeline for SV detection. Hashed arrows indicate steps that need to be taken once per reference genome. The pipeline has the steps to go from raw FASTQ reads to a finalized VCF file with annotated SV calls/genotypes across all samples in a study. Respecting Snakemake style, a template command list was written into a Snakefile, where the global rule ('all rule') set was to produce one output with all samples and SVs identified (provided in Supplementary Note 2). Conda environments were created with Python v.2.7 and v.3, given that Lumpy Smoove was written in Python v.2.7 and Snakemake was written in Python v.3 (see Supplementary Note 2).

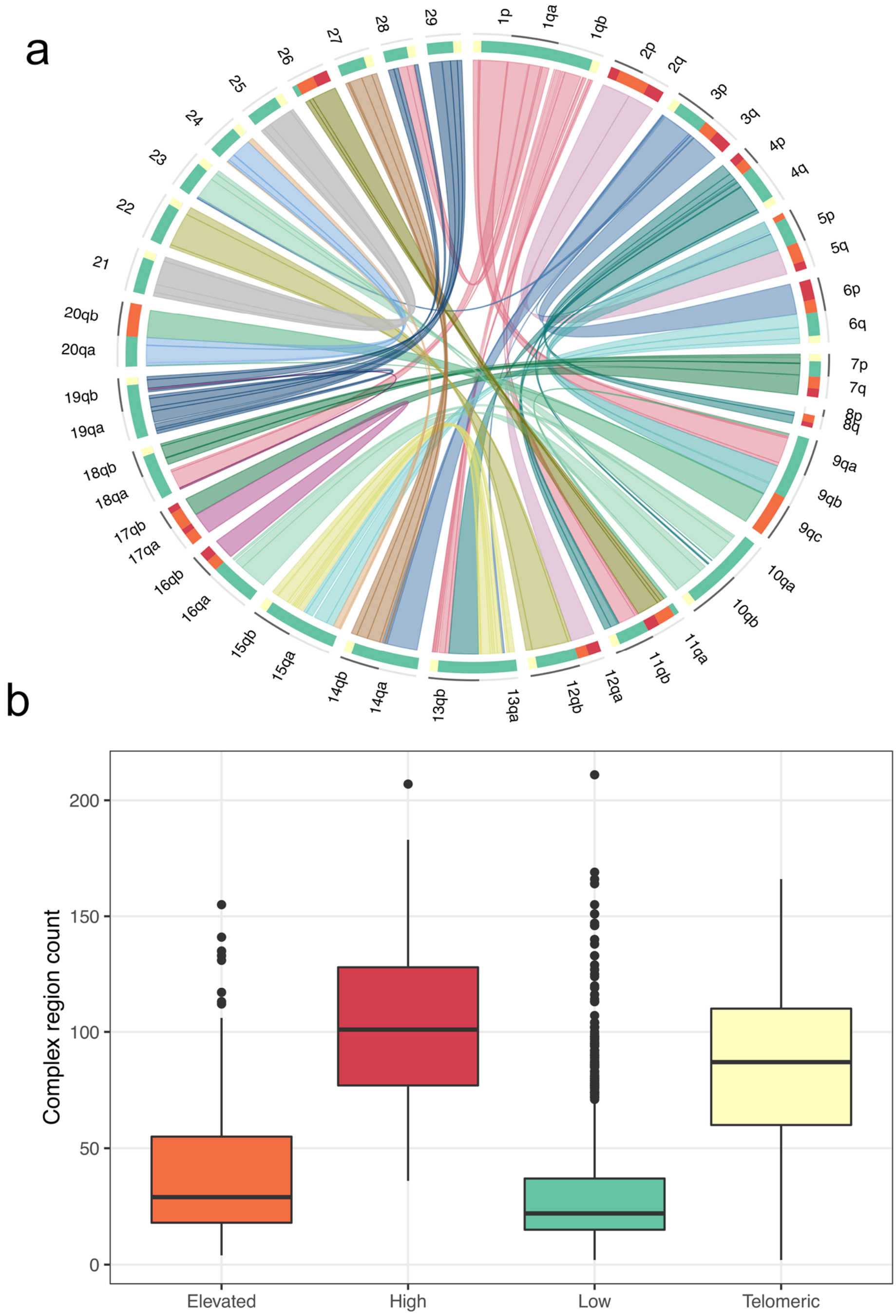


**Supplementary Figure 2**. Locations of complex regions in Atlantic salmon genome. **a** Circos plot adapted from [17], showing the 29 Atlantic salmon chromosomes divided into duplicated regions retained from the Ss4R WGD event (linked by inner circle ribbons). The main track shows each chromosome arm (names follow Atlantic salmon nomenclature [17]) divided into four categories: 'telomeres' (yellow), duplicated regions of 'low' duplicated similarity (~87% - green), duplicated regions of 'elevated' duplicated similarity (90-95% - orange) and duplicated regions of 'high' duplicated similarity (>95% - red). **b** Boxplot of counts of complex regions (defined as gap regions and regions with coverage ≥100x) across the four different categories of the genome. Complex regions were associated with an exceptionally high rate of false positive SV calls and filtered from the analysis (see Main text)

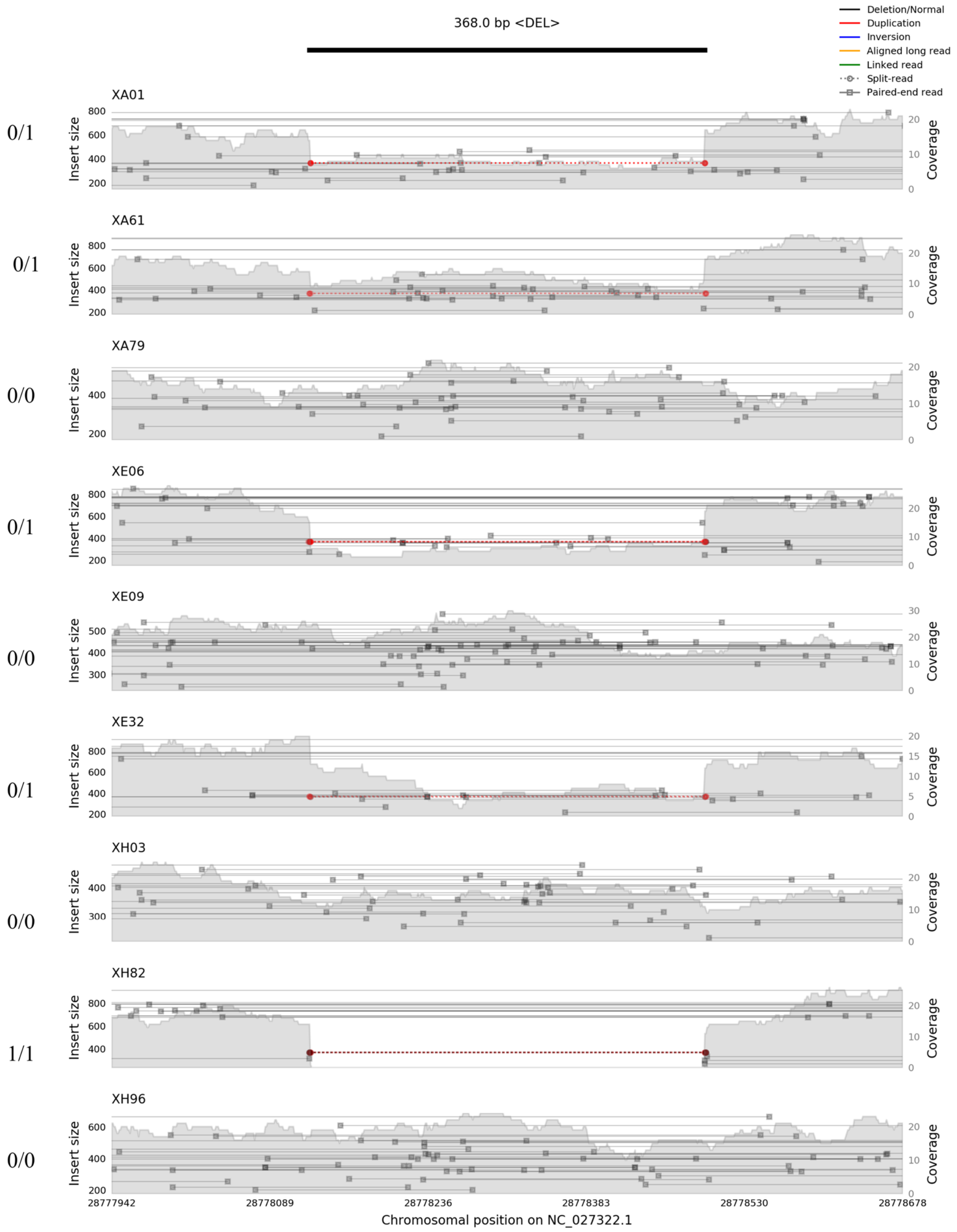


**Supplementary Figure 3.** Example of SV-plaudit image for a high confidence deletion SV call. Nine fish are shown with genotype calls given at the left of each samplot image. Also provided is the estimated breakpoint and size of the predicted deletion (black line at top of image), and a key describing the visualized evidence underlying the SV call.

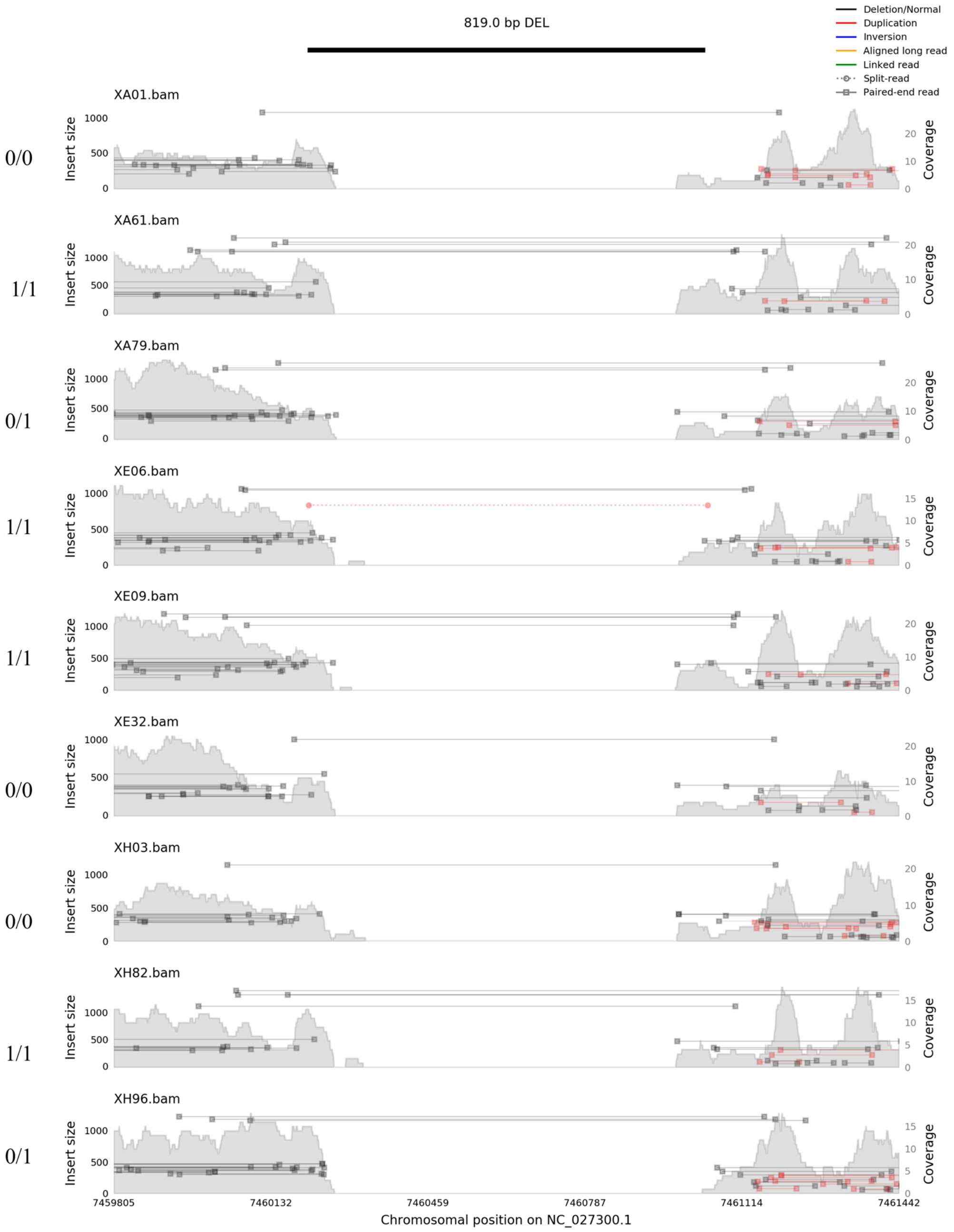


**Supplementary Figure 4.** Example of SV-plaudit image for a false positive deletion SV call excluded from further analyses. Nine fish are shown with genotype calls given at the left of each samplot image. Also provided is the estimated breakpoint and size of the predicted deletion (black line at top of image), and a key describing the visualized evidence underlying the SV call.

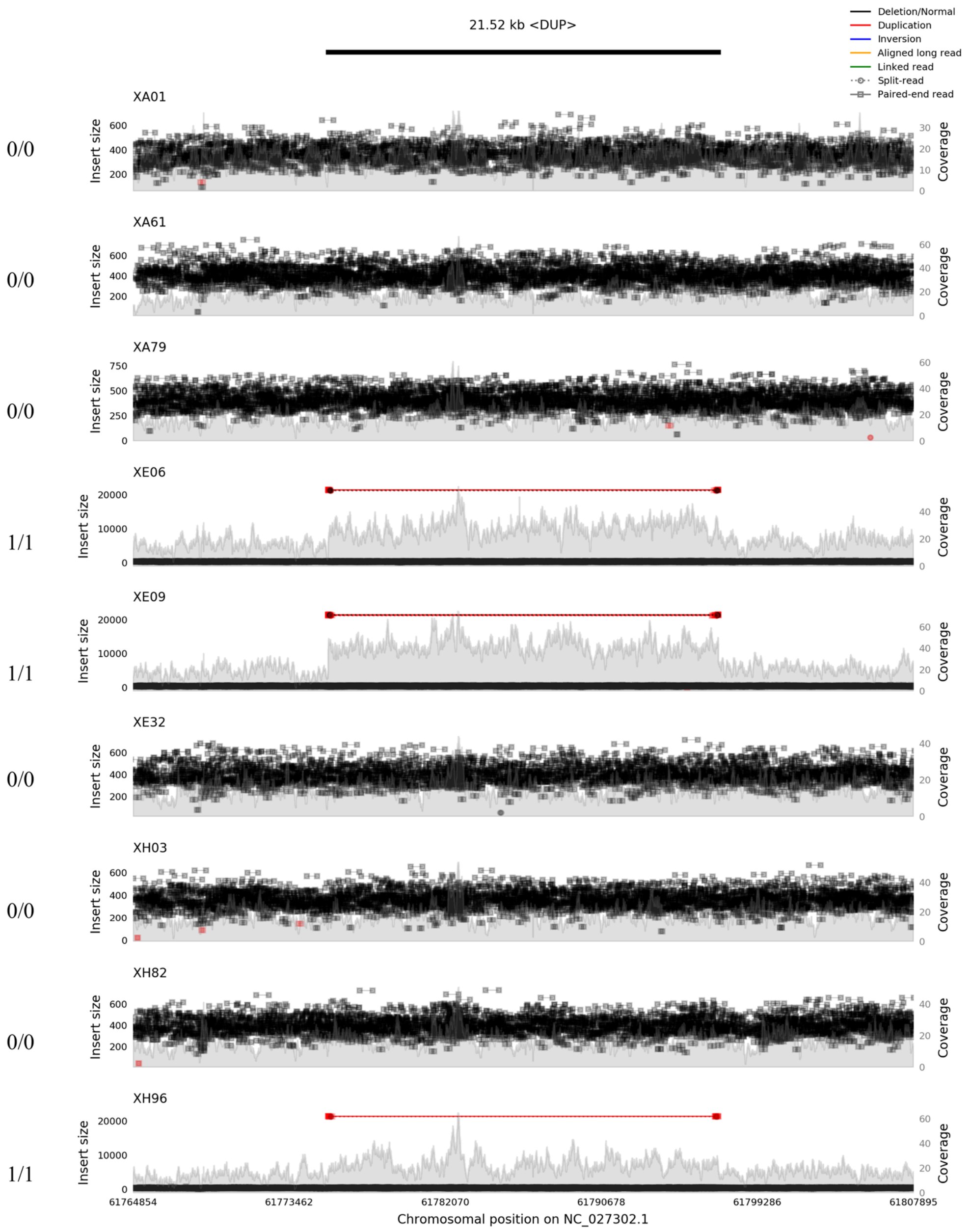


**Supplementary Figure 5.** Example of SV-plaudit image for a high confidence duplication SV call retained in further analyses. Nine fish are shown with genotype calls given at the left of each samplot image. Also provided is the estimated breakpoint and size of the predicted deletion (black line at top of image), and a key describing the visualized evidence underlying the SV call.

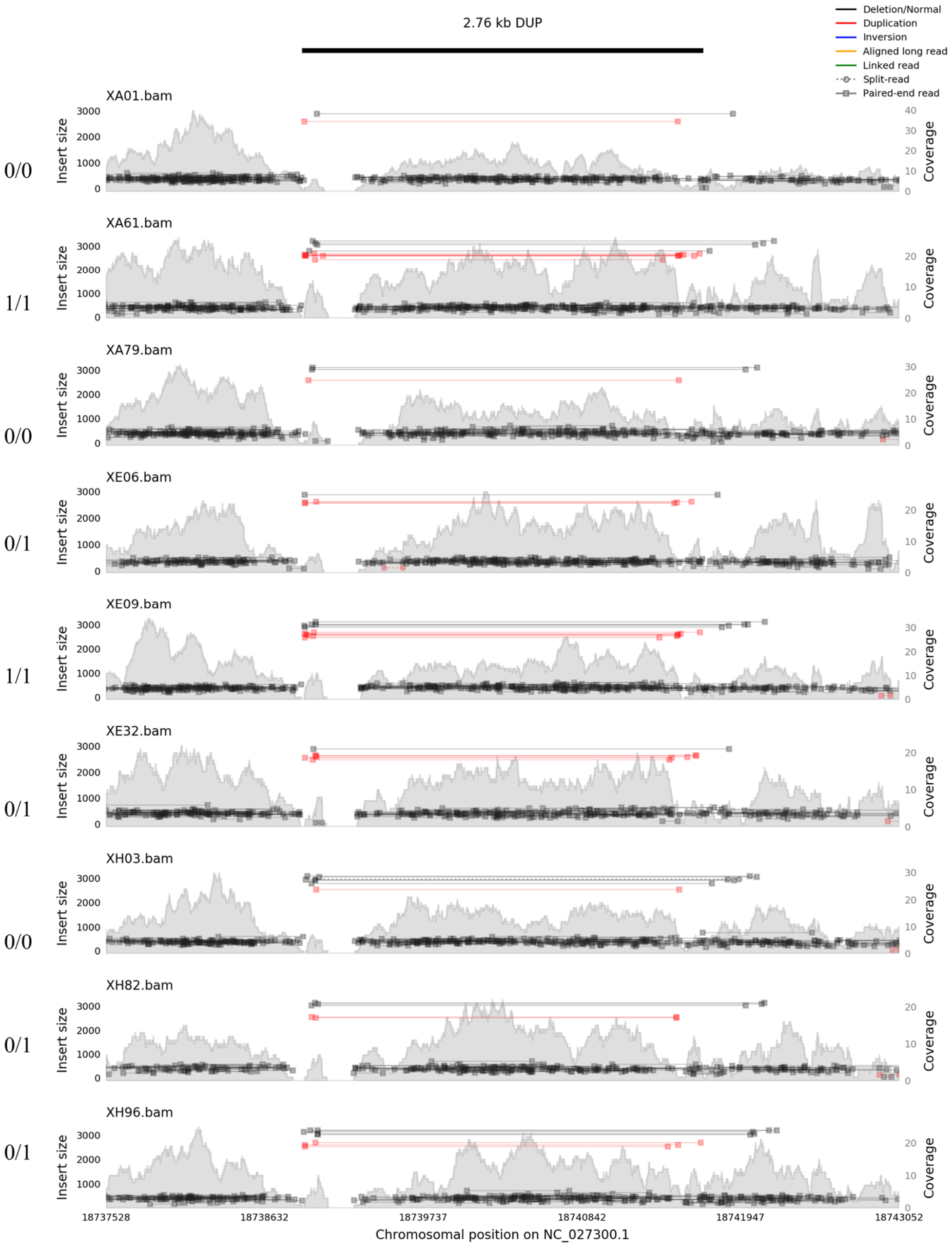


**Supplementary Figure 6.** Example of SV-plaudit image for a false positive duplication SV call excluded from further analyses. Nine fish are shown with genotype calls given at the left of each samplot image. Also provided is the estimated breakpoint and size of the predicted deletion (black line at top of image), and a key describing the visualized evidence underlying the SV call.

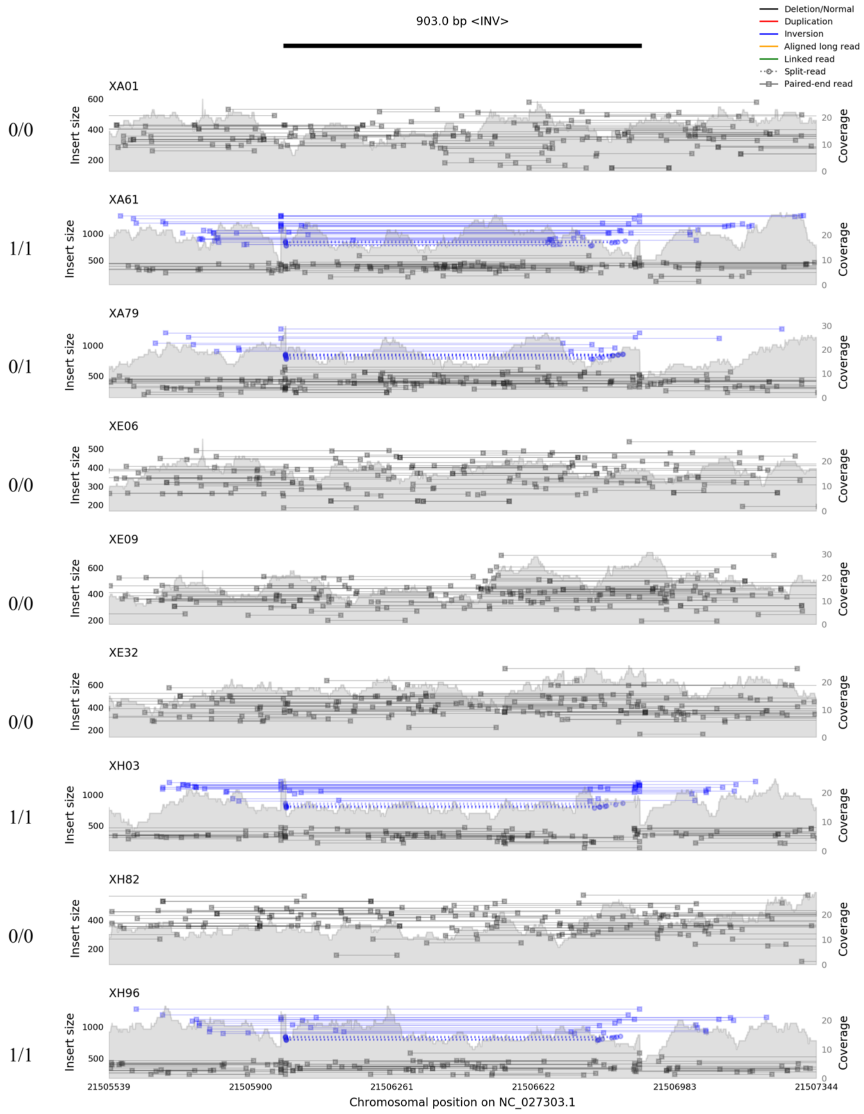


**Supplementary Figure 7.** Example of SV-plaudit image for a high confidence inversion SV call retained in further analyses. Nine fish are shown with genotype calls given at the left of each samplot image. Also provided is the estimated breakpoint and size of the predicted deletion (black line at top of image), and a key describing the visualized evidence underlying the SV call.

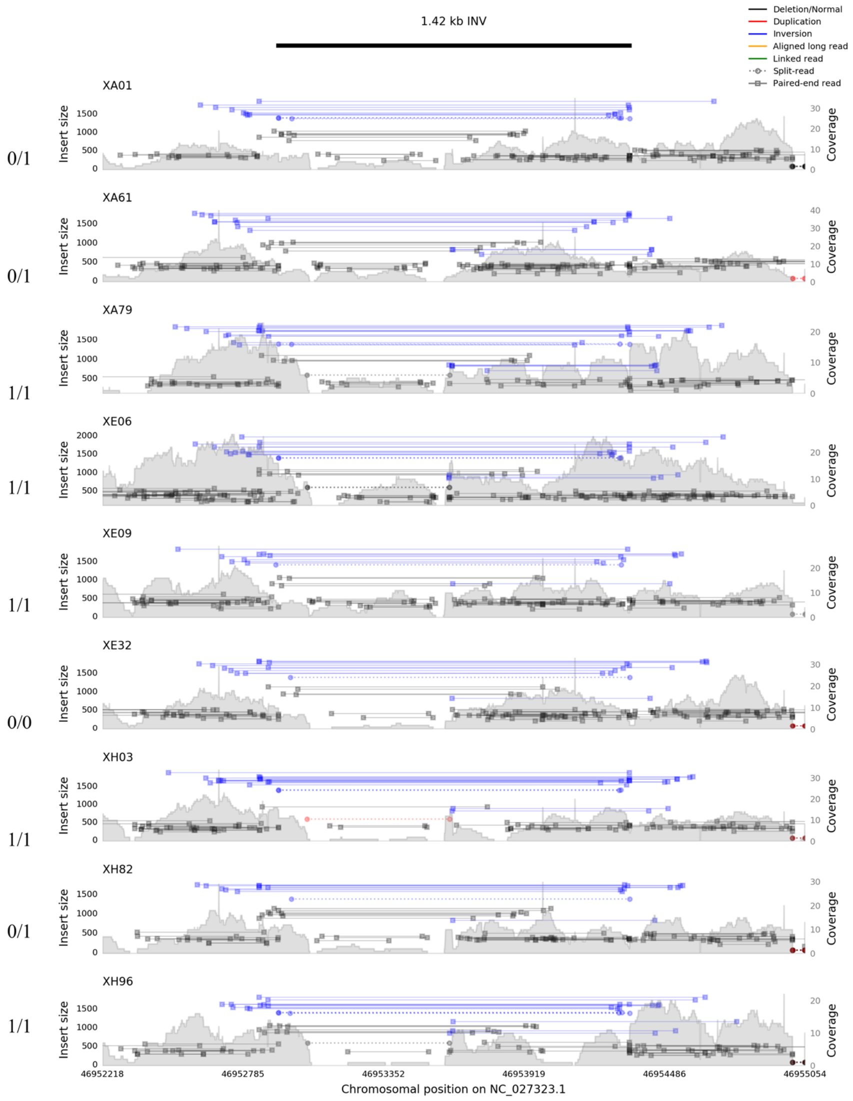


**Supplementary Figure 8.** Example of SV-plaudit image for a false positive inversion SV call excluded from further analyses. Nine fish are shown with genotype calls given at the left of each samplot image. Also provided is the estimated breakpoint and size of the predicted deletion (black line at top of image), and a key describing the visualized evidence underlying the SV call.

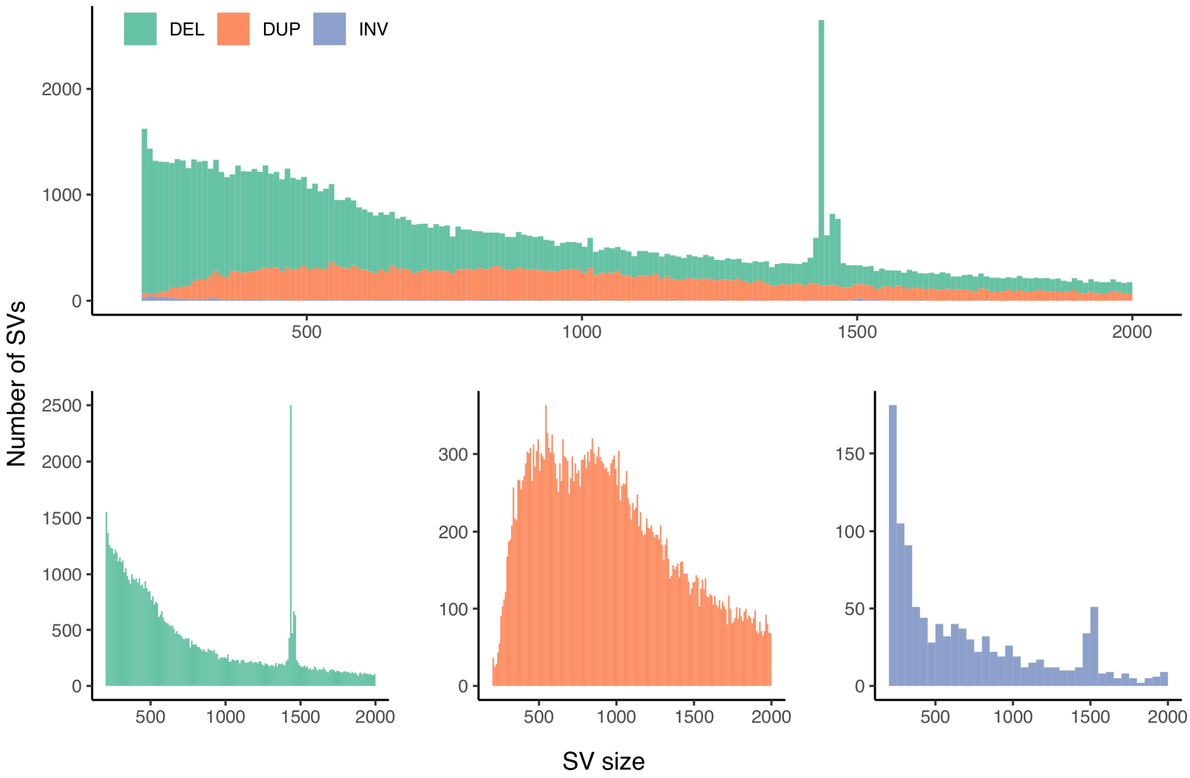


**Supplementary Figure 9**. SV sizes before filtering and SV-plaudit curation. Note the peak observed in the deletion calls, which remained after SV-plaudit curation, and is further characterized in Fig. 2.

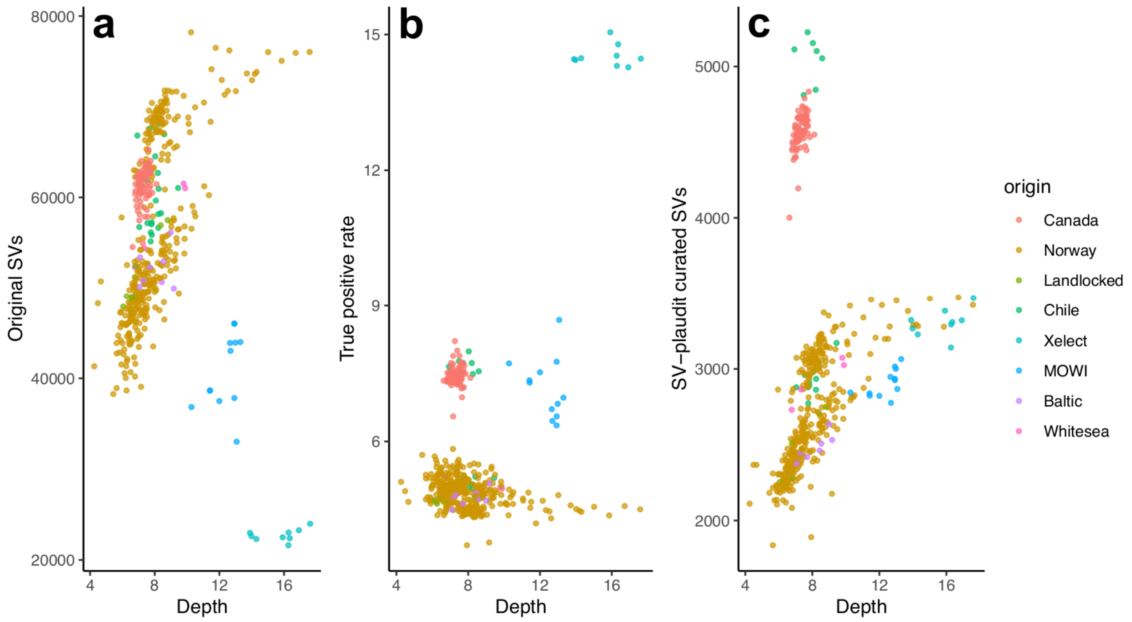


**Supplementary Figure 10.** Sequencing depth was not a strong predictor of the final number of high-confidence SVs retained after SV-plaudit curation. The scatterplots show sequencing depth plotted against **a** the number of original/unfiltered SV calls, **b** the true positive rate (%) for SV calls (number of retained high confidence SV calls after SV-plaudit validation / number of original SV calls) and **c** the final number of SV-plaudit curated SV calls. The origin of each sample is shown by a different colour according to the given key (see Supplementary Table 1). Our hypothesis was that increased sequencing depth would lead to a lower FDR, along with an increased true positive rate. However, there was no significant correlation between sequencing depth and the number of unfiltered SVs (part **a**, Pearson's correlation coefficient = 0.07, $P$ = 0.11) and a weak but significant positive correlation between sequencing depth and true positive SV call rate (part **b**; Pearson's correlation coefficient = 0.34, $P$ < 0.001). Observing the final SV-plaudit curated data (part **c**), it can be seen that a few samples skew the expected relationship, i.e. either have relatively high sequencing depth with a relatively high number of unfiltered SVs and relatively low true positive SV call rate. It can also be seen that most the variability is explained by study differences; the data was derived from five different studies, which include variation not just in sequencing depth, but also biology (i.e. distinct geographical locations, farmed vs. wild fish), as well as technological factors (four distinct sequencing platforms; many different labs involved in DNA extraction/library preps). Together, these features may obscure the expected relationship between SV detection and sequencing depth.

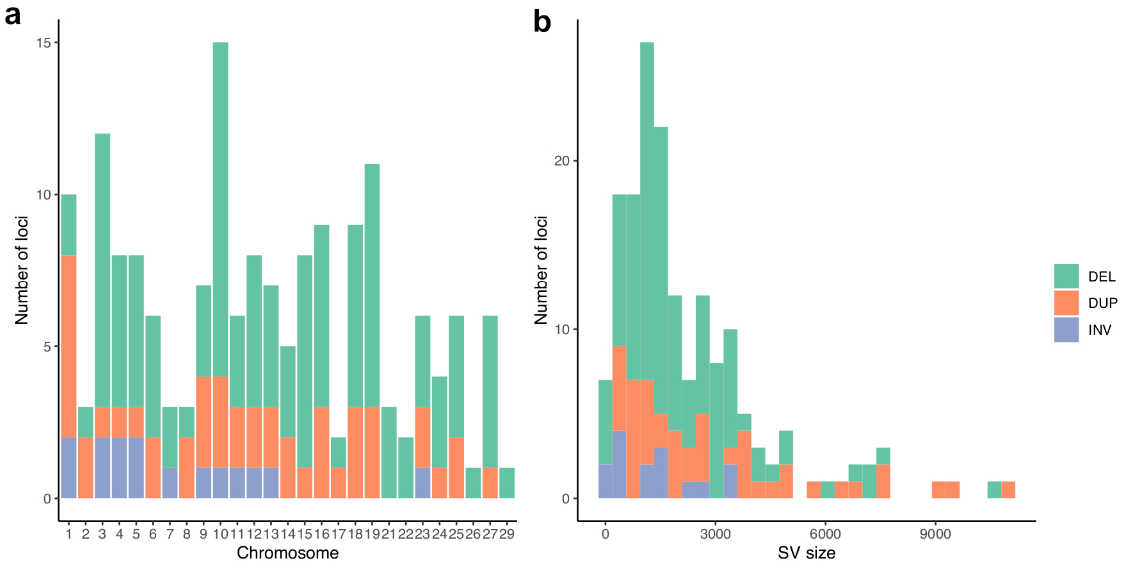


**Supplementary Figure 11.** Summary of 168 SV regions used for MinION amplicon sequencing to validate Lumpy/SVtyper SV and genotype calls. **a** Number of SV regions targeted on different salmon chromosomes. **b** Size distribution of SV regions targeted by PCR.

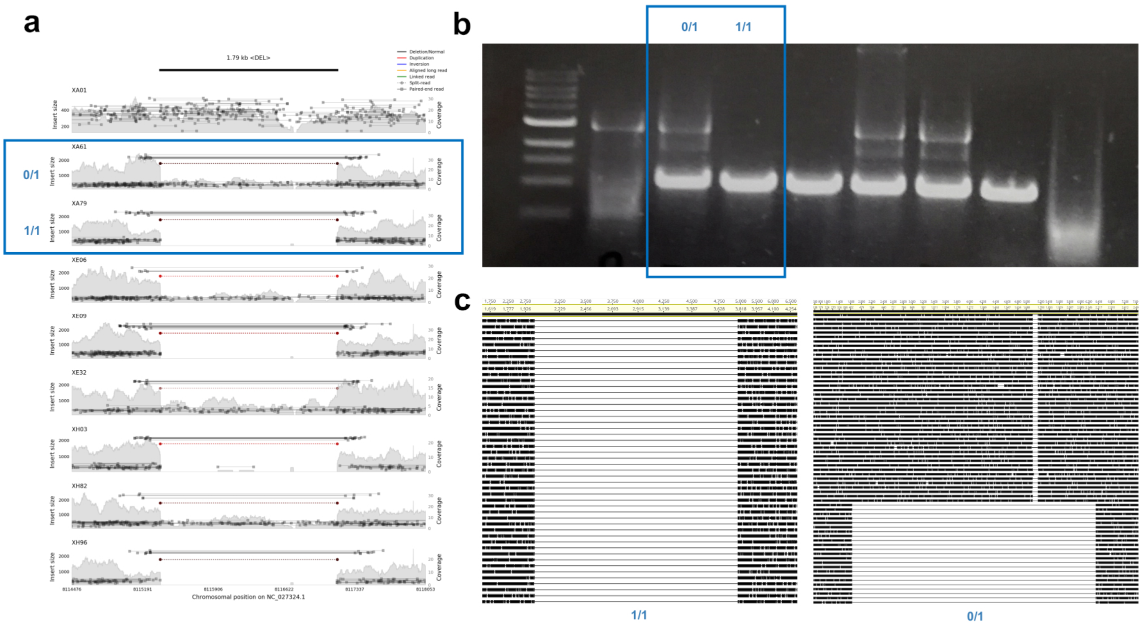


**Supplementary Figure 12**. Example of congruence between SV/genotype calls and data generated by MinION amplicon sequencing. **a** SV-plaudit deletion plot for 9 tested salmon individuals. Circled in blue are one 0/1 genotype call and one 1/1 genotype call. **b** agarose gel image for the same individuals, with two calls highlighted in the gel image, showing the same genotypes. **c** Alignments of amplicons sequenced for the samples highlighted in part **b**.

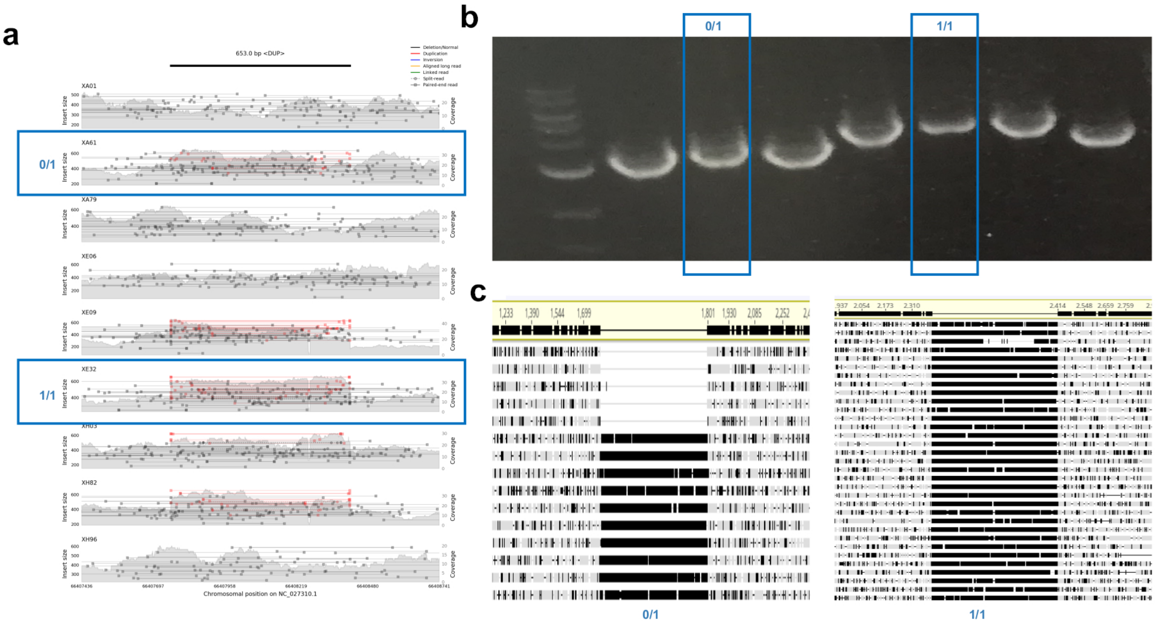


**Supplementary Figure 13.** Example of congruence between SV/genotype calls and data generated by MinION amplicon sequencing. **a** SV-plaudit duplication plot for 9 tested salmon individuals. Circled in blue are one 0/1 genotype call and one 1/1 genotype call. **b** Agarose gel image for the same individuals, with two calls highlighted in the gel image, showing the same genotypes. **c** Alignments of amplicons sequenced for the same samples highlighted in **b** .

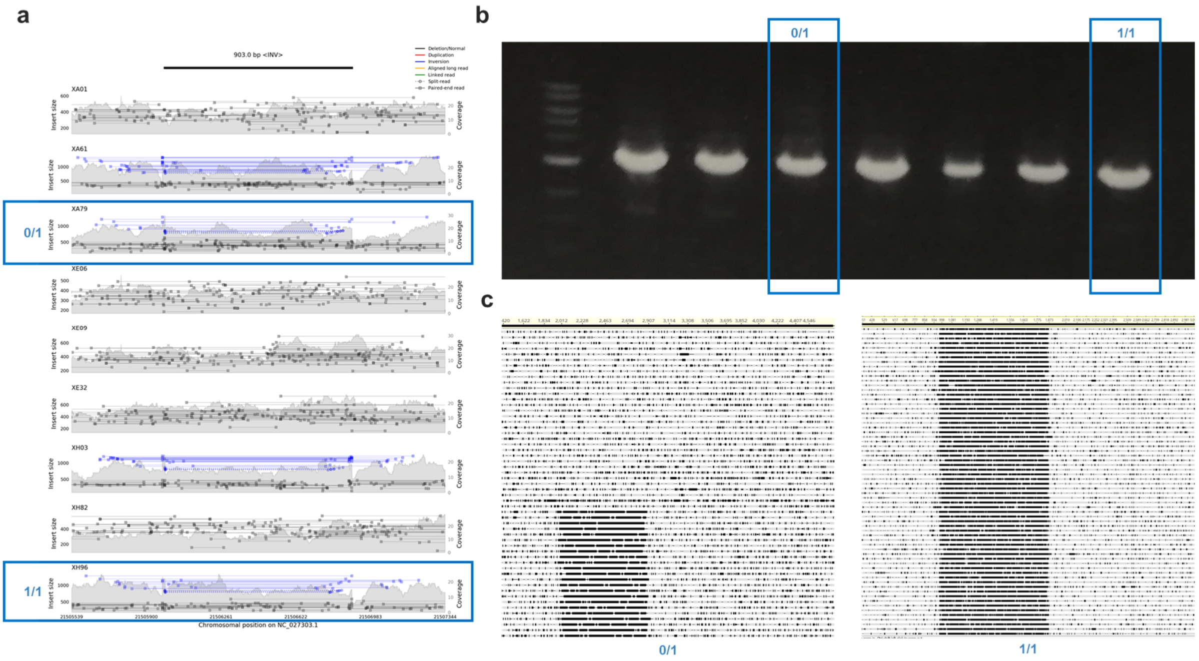


**Supplementary Figure 14**. Example of congruence between SV/genotype calls and data generated by MinION amplicon sequencing. **a** SV-plaudit inversion plot for 9 tested salmon individuals. Circled in blue are one 0/1 genotype call and one 1/1 genotype call. **b** Agarose gel image for the same individuals, with two calls highlighted in the gel image, showing the same genotypes. **c** Alignments of amplicons sequenced for the same samples highlighted in **b**.

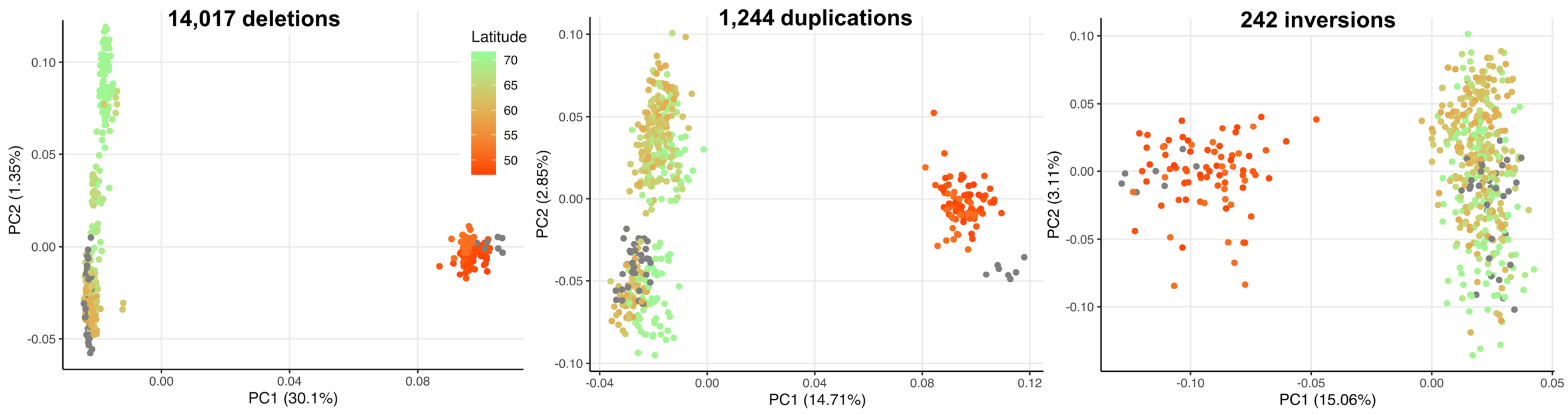


**Supplementary Figure 15.** PCAs showing the same data presented in Fig. 1g-i (main text), except visualized according to latitude. For all SV types, PC1 separates European and Canadian salmon (orange symbols; wild Canadian fish were all sampled from lower latitudes than all wild European salmon). For the deletion genotypes, PC2 separates populations according to latitude, which is expected (see main text). The grey symbols depict farmed salmon individuals.

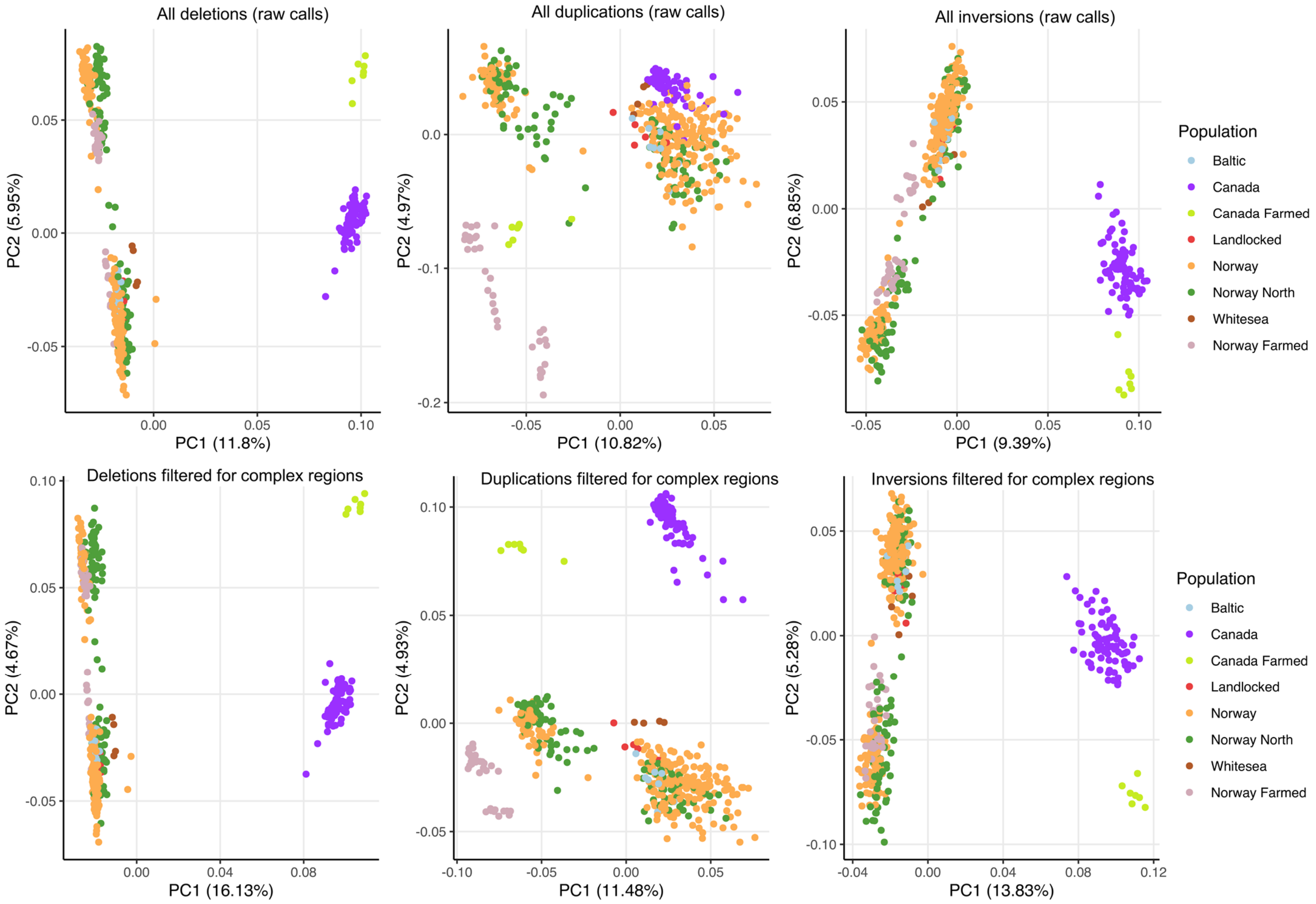


**Supplementary Figure 16.** PCA analyses done on SV genotype calls prior to SV-plaudit curation for i) the complete set of unfiltered SV genotype calls prior to filtering complex regions or SV-plaudit curation (top panel) and ii) the subset of SV genotype calls remaining after filtering complex regions in the genome, but prior to SV-plaudit curation (bottom panel). Consistent with the filtering of unreliable SV calls during SV-plaudit curation, both sets fail to capture expected population genetic structure at high resolution. For instance, in both datasets, duplications fail to separate Canadian and European salmon. Further, both datasets fail to split European salmon by latitude for all SV types (see Supplementary Figure 15) and erroneously place farmed salmon populations. For references supporting these deductions, see main text.

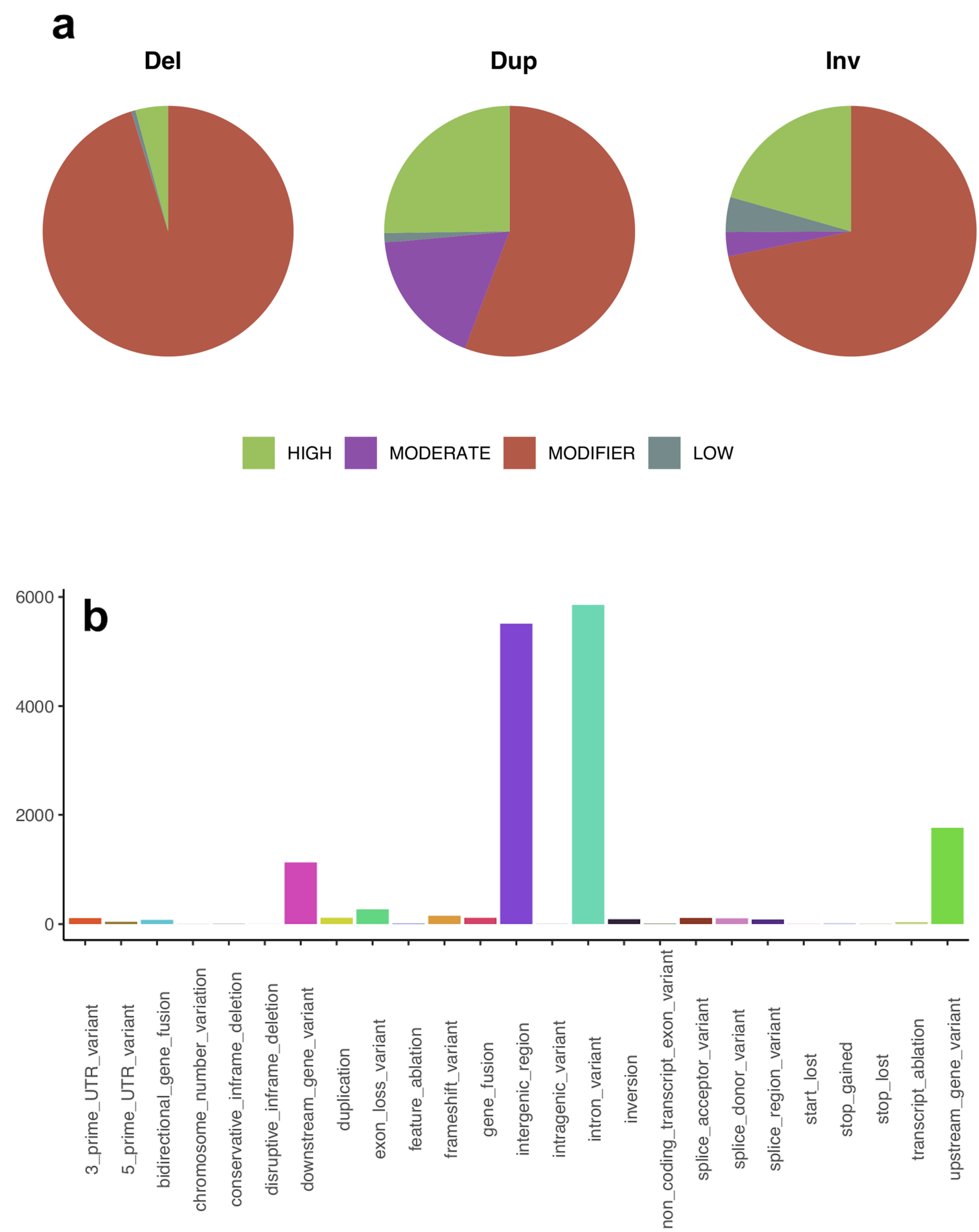


**Supplementary Figure 17.** SV annotation by SnpEff **a** Proportion of putative impacts predicted by snpEff for each SV type shown for all high-confidence Atlantic salmon SVs. **b** Number of SVs for each effect as annotated by snpEff.

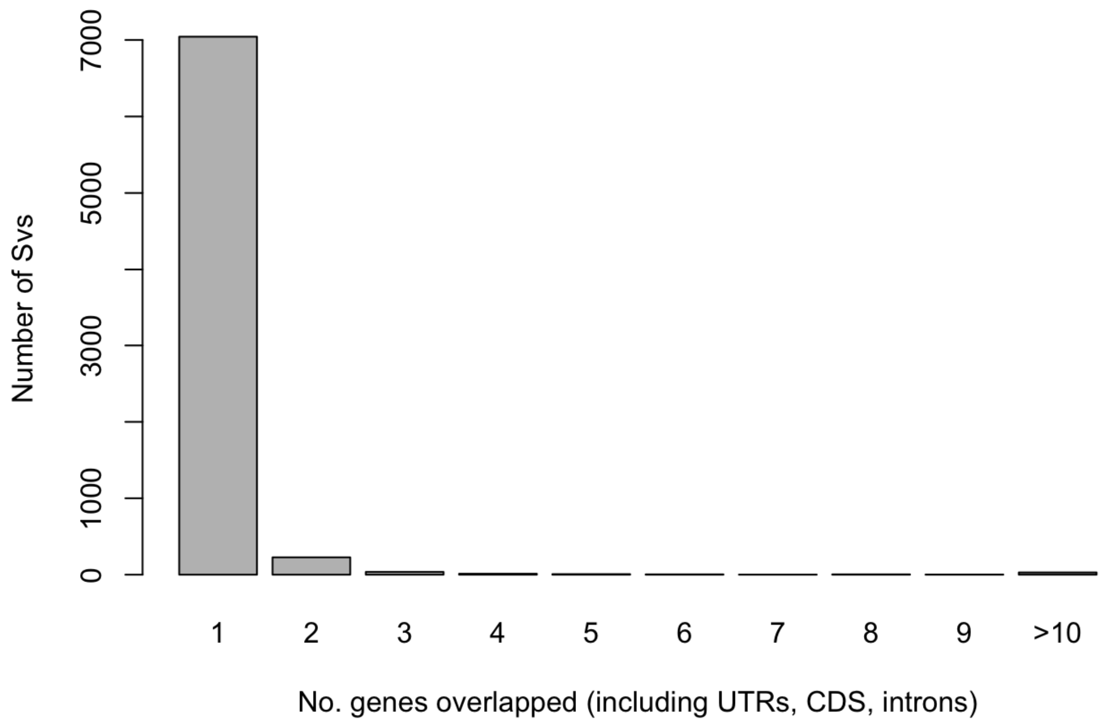


**Supplementary Figure 18**. Overlap between high confidence SVs and protein coding genes in the ICSASG_v2 annotation.

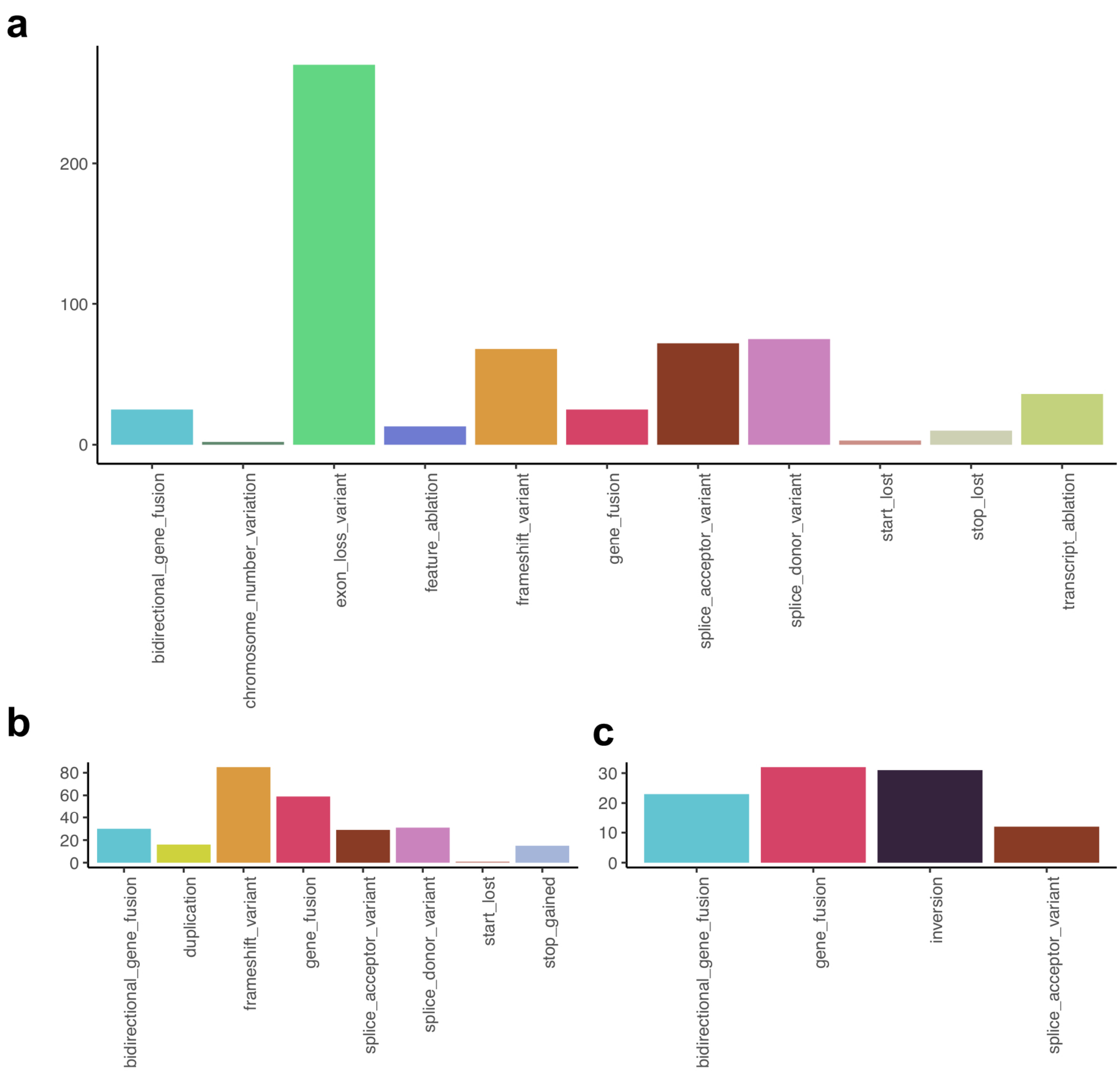


**Supplementary Figure 19.** Number of high impact annotations per snpEff effect for high-confidence Atlantic salmon SVs. **a** deletions, **b** duplications, **c** inversions.

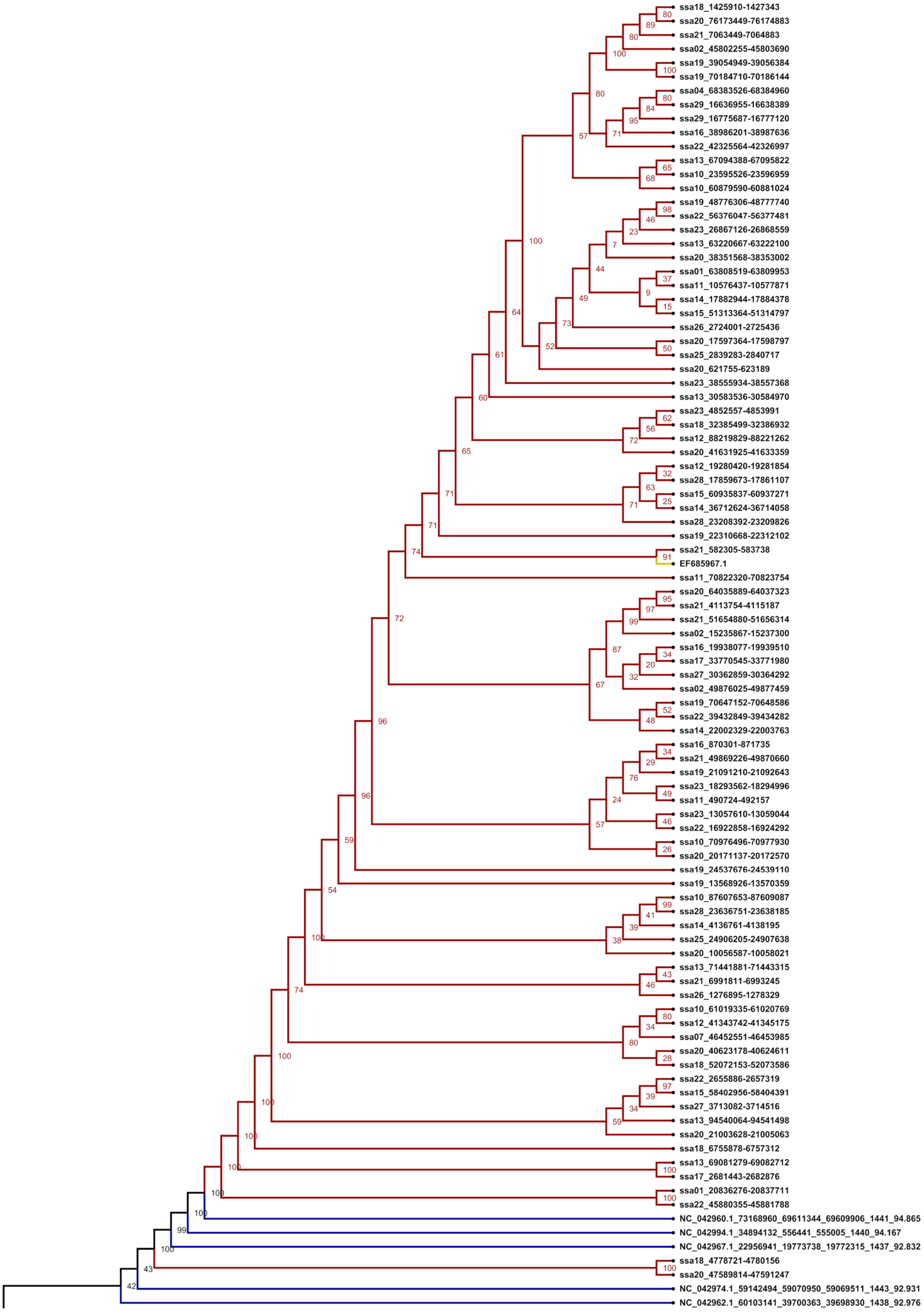


Cont. on next page

Cont. from last page

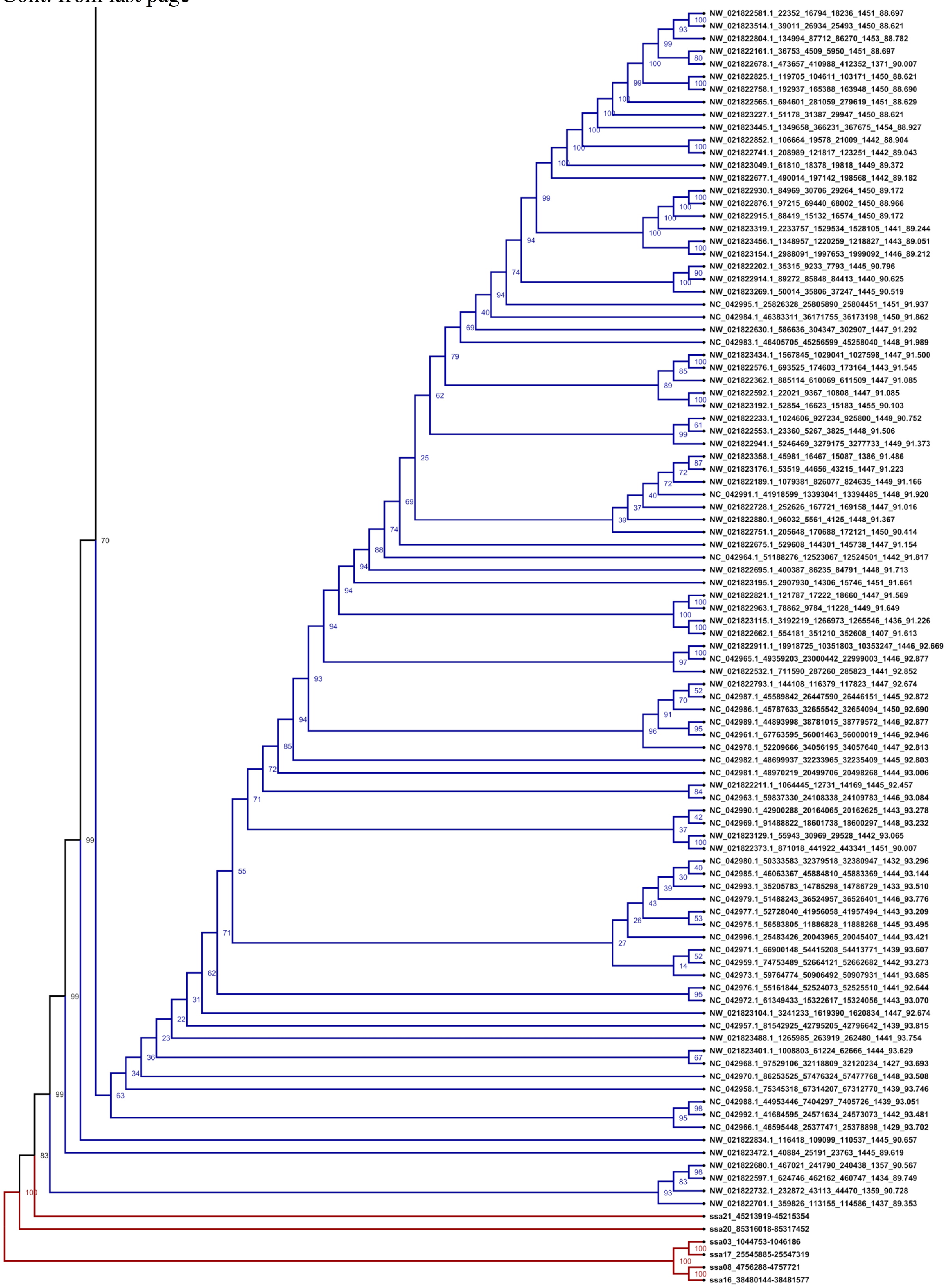


**Supplementary Figure 20.** Maximum likelihood tree presented in Main Text Fig. 2 including sample identifiers, genomic locations of pTSsa2 sequences and bootstrap values**.**

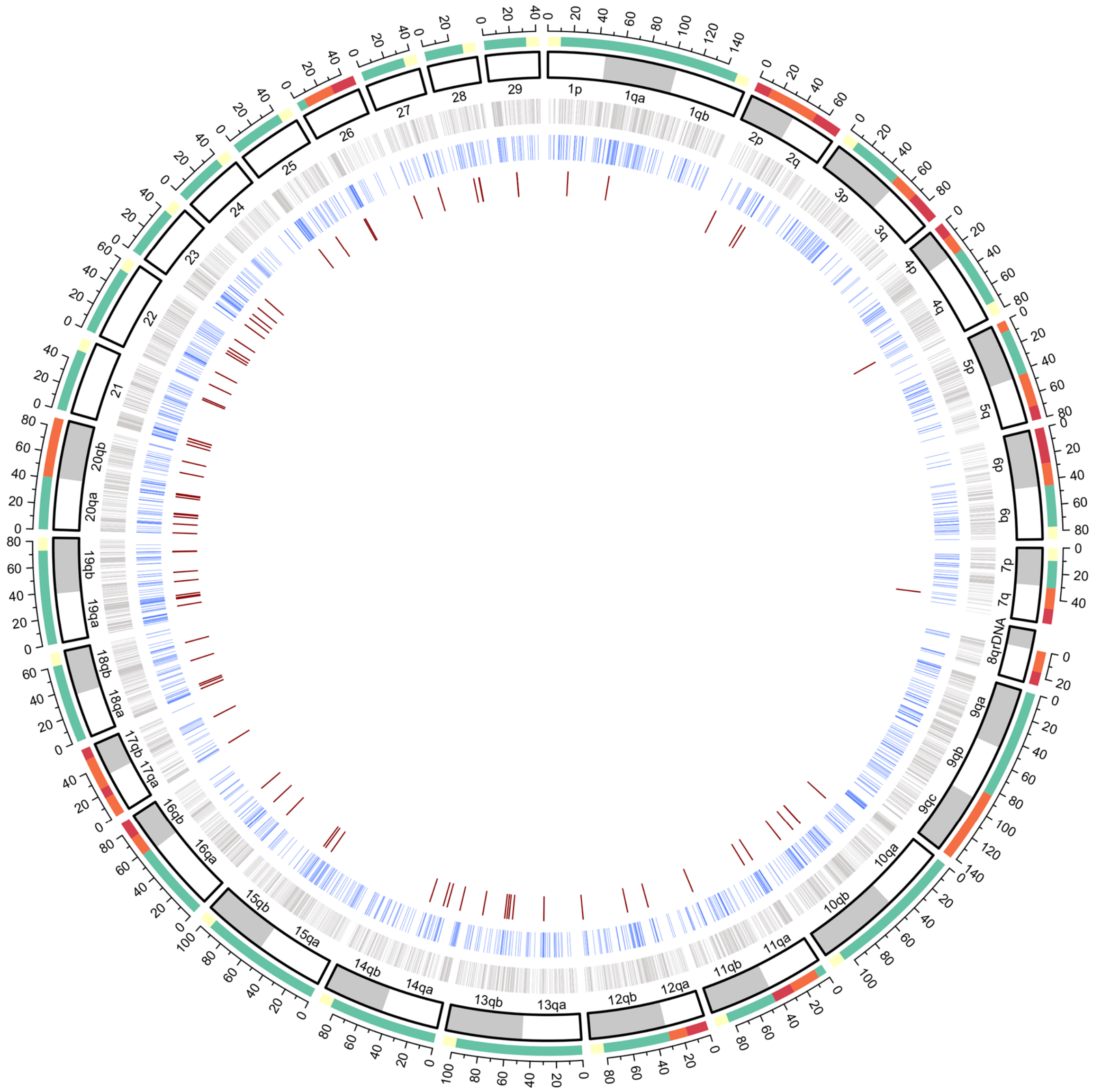


**Supplementary Figure 21.** Circos plot showing the genomic locations of pTSsa2 sequences in the Atlantic salmon genome for i) 94 polymorphic deletion sequences used in the tree shown in Fig. 2 (red lines, internal track), ii) 1,519 sequences within 1432-1436bp deletion peak before SV-plaudit filtering, sharing ≥95% and ≥ 98% coverage BLAST identity and coverage to the reference pTSsa2 sequence (blue lines, middle track) and iii) all pTSsa2 in the unmasked Atlantic salmon genome sharing ≥95% and ≥ 98% coverage BLAST identity to the reference pTSsa2 (grey line, outer track). The colours on the outside of chromosome arms, which are depicted with established *S. salar* nomenclature[17], depict the regions highlighted in Fig. 1b and Supplementary Fig. 2.

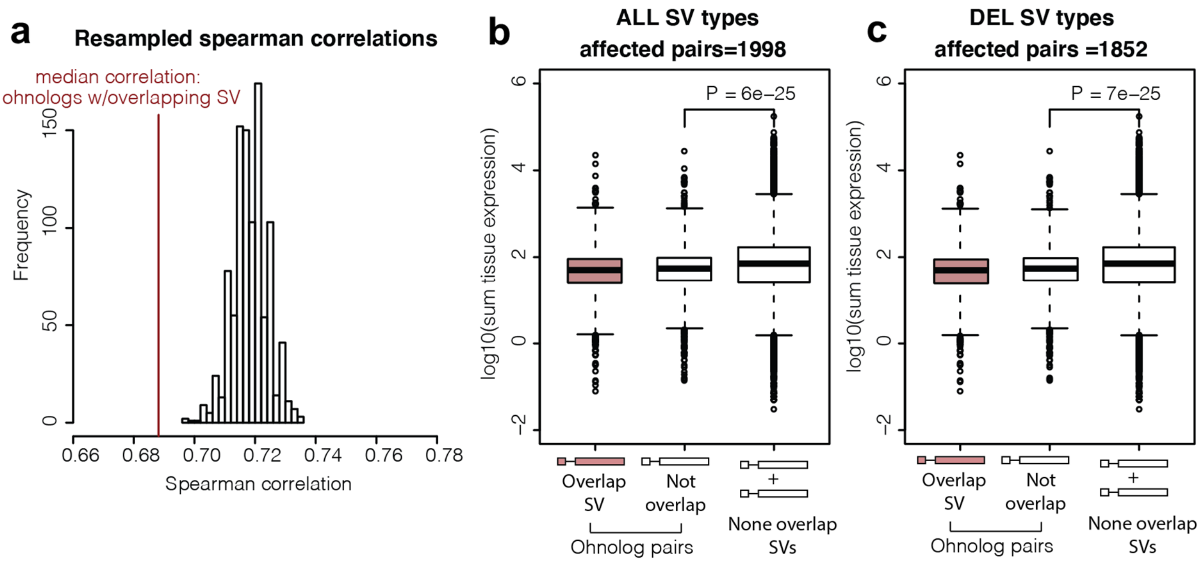


**Supplementary Figure 22.** Expression characteristics of ohnologs depending on SV overlap. **a** Spearman's correlation between Ss4R ohnolog pairs where one gene is overlapped by an SV in comparison to randomly resampled ohnolog pairs that do not overlap SVs. Also shown, both for all SVs (**a**) and just deletions (**b**), is the expression level of ohnologs (the sum of expression across 15 tissues), with different boxplots representing ohnologs overlapped by an SV (left) vs. the other gene in the same ohnolog pair (that does not overlap an SV) (middle) and ohnolog pairs where neither gene is overlapped by any SV.

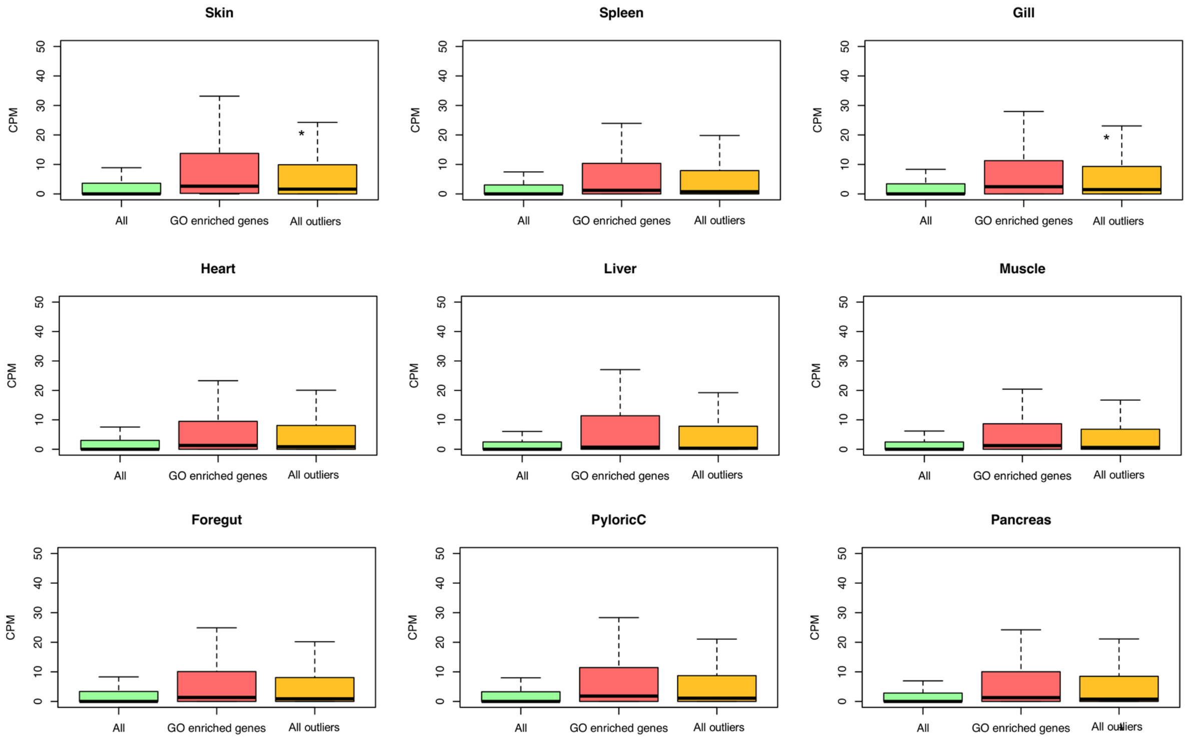


**Supplementary Figure 23.** Tissue expression levels (counts per million, CPM) comparing 44,469 genes in a transcriptome ('All') with genes linked by SnpEff to the 584 SV outliers ('All outliers') and the subset of 326 genes contributing to enriched biological processes by GO analysis ('GO enriched genes'). The asterisk (*) indicates a *P* value <0.05. Brain is shown in Fig. 4d (main text).

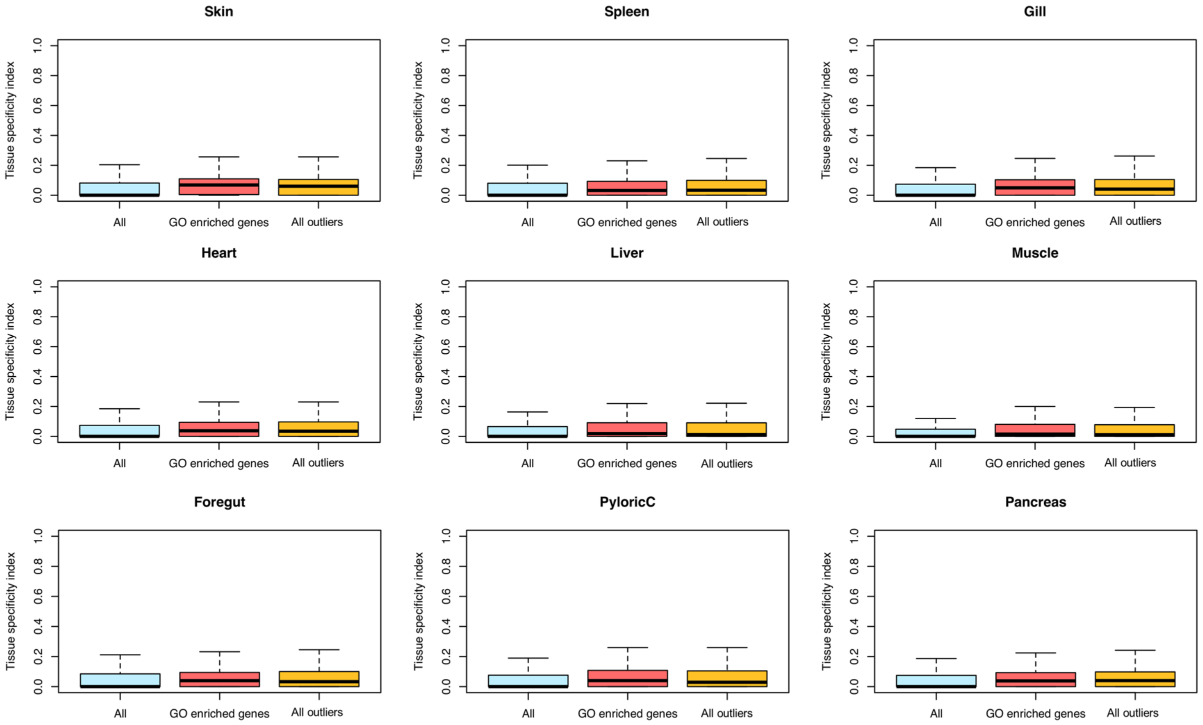


**Supplementary Figure 24.** Tissue specificity (see Methods) comparing 44,469 genes in a transcriptome ('All') with genes linked by SnpEff to the 584 SV outliers ('All outliers') and the subset of 326 genes contributing to enriched biological processes by GO analysis ('GO enriched genes'). Brain is shown in Fig. 4d (main text).

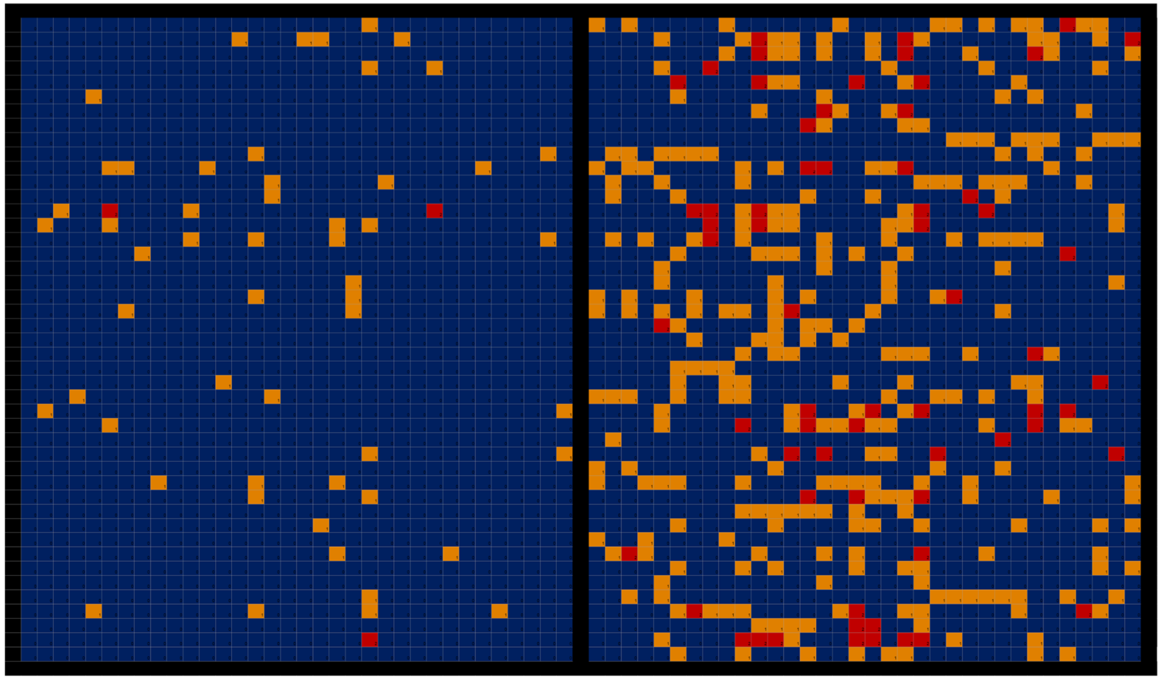


| Chr | Start | End | Unique ID | Freq 0/0 Wild | Freq 0/1 Wild | Freq 1/1 Wild | Freq 0/0 Farmed | Freq 0/1 Farmed | Freq 1/1 Farmed |
|---|---|---|---|---|---|---|---|---|---|
| ssa01 | 53229610 | 54031283 | Outlier-25 | 0.96 | 0.04 | 0.00 | 0.65 | 0.32 | 0.03 |
| ssa01 | 131617069 | 131617450 | Outlier-54 | 0.92 | 0.07 | 0.01 | 0.62 | 0.29 | 0.09 |
| ssa01 | 132558562 | 132558973 | Outlier-55 | 0.98 | 0.02 | 0.00 | 0.68 | 0.24 | 0.09 |
| ssa02 | 55721269 | 55721405 | Outlier-87 | 0.99 | 0.01 | 0.00 | 0.85 | 0.12 | 0.03 |
| ssa03 | 18151575 | 18151741 | Outlier-96 | 1.00 | 0.00 | 0.00 | 0.76 | 0.12 | 0.12 |
| ssa03 | 21969738 | 21969900 | Outlier-98 | 1.00 | 0.00 | 0.00 | 0.88 | 0.12 | 0.00 |
| ssa03 | 33657659 | 33657805 | Outlier-101 | 0.97 | 0.02 | 0.01 | 0.82 | 0.12 | 0.06 |
| ssa04 | 45249290 | 45250396 | Outlier-135 | 1.00 | 0.00 | 0.00 | 0.88 | 0.09 | 0.03 |
| ssa04 | 47954778 | 47954884 | Outlier-136 | 0.98 | 0.02 | 0.00 | 0.74 | 0.26 | 0.00 |
| ssa04 | 75012242 | 75012417 | Outlier-142 | 0.96 | 0.04 | 0.00 | 0.74 | 0.26 | 0.00 |
| ssa05 | 17288655 | 17291404 | Outlier-148 | 0.91 | 0.09 | 0.00 | 0.68 | 0.24 | 0.09 |
| ssa05 | 41568467 | 41572967 | Outlier-158 | 0.96 | 0.04 | 0.00 | 0.71 | 0.29 | 0.00 |
| ssa09 | 80636406 | 80636651 | Outlier-225 | 0.98 | 0.02 | 0.00 | 0.82 | 0.15 | 0.03 |
| ssa09 | 102103011 | 102103264 | Outlier-226 | 0.92 | 0.07 | 0.02 | 0.71 | 0.15 | 0.15 |
| ssa09 | 103850700 | 103851015 | Outlier-234 | 0.94 | 0.06 | 0.00 | 0.74 | 0.18 | 0.09 |
| ssa09 | 139513277 | 139513675 | Outlier-243 | 0.93 | 0.07 | 0.00 | 0.65 | 0.32 | 0.03 |
| ssa10 | 53986929 | 53988379 | Outlier-255 | 0.97 | 0.03 | 0.00 | 0.76 | 0.21 | 0.03 |
| ssa11 | 10118880 | 10123943 | Outlier-278 | 1.00 | 0.00 | 0.00 | 0.88 | 0.12 | 0.00 |
| ssa11 | 61339068 | 61340955 | Outlier-288 | 0.99 | 0.01 | 0.00 | 0.88 | 0.12 | 0.00 |
| ssa12 | 33326446 | 33326756 | Outlier-299 | 0.96 | 0.04 | 0.00 | 0.76 | 0.21 | 0.03 |
| ssa12 | 67331886 | 67333727 | Outlier-303 | 0.92 | 0.07 | 0.00 | 0.71 | 0.26 | 0.03 |
| ssa13 | 2636866 | 2636988 | Outlier-307 | 0.98 | 0.02 | 0.00 | 0.82 | 0.15 | 0.03 |
| ssa13 | 9914689 | 9924395 | Outlier-309 | 1.00 | 0.00 | 0.00 | 0.85 | 0.15 | 0.00 |
| ssa13 | 53921156 | 53921275 | Outlier-320 | 0.95 | 0.04 | 0.00 | 0.74 | 0.24 | 0.03 |
| ssa13 | 68700203 | 68721406 | Outlier-325 | 0.99 | 0.01 | 0.00 | 0.88 | 0.12 | 0.00 |
| ssa14 | 10890695 | 10890938 | Outlier-347 | 0.99 | 0.01 | 0.00 | 0.76 | 0.21 | 0.03 |
| ssa14 | 13696392 | 13696763 | Outlier-349 | 0.94 | 0.06 | 0.00 | 0.68 | 0.32 | 0.00 |
| ssa14 | 28688223 | 28688473 | Outlier-360 | 0.91 | 0.06 | 0.03 | 0.74 | 0.12 | 0.15 |
| ssa14 | 71904738 | 71959507 | Outlier-370 | 0.93 | 0.04 | 0.02 | 0.62 | 0.26 | 0.12 |
| ssa18 | 15447276 | 15447639 | Outlier-399 | 1.00 | 0.00 | 0.00 | 0.94 | 0.03 | 0.03 |
| ssa18 | 15665311 | 15665588 | Outlier-414 | 0.94 | 0.06 | 0.00 | 0.79 | 0.09 | 0.12 |
| ssa18 | 24680046 | 24680231 | Outlier-415 | 0.99 | 0.01 | 0.00 | 0.85 | 0.15 | 0.00 |
| ssa18 | 46500281 | 47195996 | Outlier-419 | 0.93 | 0.07 | 0.00 | 0.65 | 0.35 | 0.00 |
| ssa19 | 2109421 | 2123074 | Outlier-424 | 0.97 | 0.03 | 0.00 | 0.74 | 0.18 | 0.09 |
| ssa19 | 23835619 | 23836118 | Outlier-444 | 0.99 | 0.01 | 0.00 | 0.76 | 0.24 | 0.00 |
| ssa19 | 31446481 | 31447749 | Outlier-449 | 1.00 | 0.00 | 0.00 | 0.76 | 0.24 | 0.00 |
| ssa19 | 72792415 | 72792564 | Outlier-474 | 1.00 | 0.00 | 0.00 | 0.91 | 0.09 | 0.00 |
| ssa22 | 16640863 | 16643241 | Outlier-506 | 0.94 | 0.06 | 0.00 | 0.74 | 0.21 | 0.06 |
| ssa25 | 13389967 | 13390308 | Outlier-544 | 0.99 | 0.01 | 0.00 | 0.76 | 0.24 | 0.00 |
| ssa25 | 37644994 | 37645344 | Outlier-551 | 1.00 | 0.00 | 0.00 | 0.91 | 0.09 | 0.00 |
| ssa26 | 14396169 | 14397980 | Outlier-559 | 0.93 | 0.07 | 0.00 | 0.71 | 0.29 | 0.00 |
| ssa28 | 3675867 | 3680686 | Outlier-569 | 0.90 | 0.09 | 0.01 | 0.65 | 0.26 | 0.09 |
| ssa28 | 8847491 | 8847963 | Outlier-579 | 0.99 | 0.01 | 0.00 | 0.79 | 0.15 | 0.06 |
| ssa28 | 23454925 | 23455114 | Outlier-589 | 0.92 | 0.04 | 0.04 | 0.68 | 0.12 | 0.21 |
| ssa29 | 22608376 | 22608582 | Outlier-594 | 0.96 | 0.04 | 0.00 | 0.76 | 0.24 | 0.00 |

**Supplementary Figure 25**. Heatmap showing individual SV genotypes for 45 SV outliers linked to synapse genes in the salmon genome. SVs have been filtered to only include rare SVs in wild fish. The blue, yellow and red squares depict homozygous 0/0 (lacking SV), heterozygous 0/1 (one SV copy), and homozygous 1/1 (two SV copies) genotypes, respectively. Each column represents a different fish individual. The SVs are organized from top to bottom with respect to their location in the genome.

**Supplementary Note 1**. Python script used to extract gap regions in the ICSASG_v2 genome and convert the output to a BED file.

Obtained from: https://www.danielecook.com/generate-a-bedfile-of-masked-ranges-a-fasta-file/

Usage:

```
python generate_masked_ranges.py <fasta_file> > output_ranges.txt
```

Code:

```
#!bin/python

import gzip
import io
import sys
import os

# This file will generate a bedfile of the masked regions a fasta file.

# STDIN or arguments
if len(sys.argv) > 1:

    # Check file type
    if sys.argv[1].endswith(".fa.gz"):
        input_fasta = io.TextIOWrapper(io.BufferedReader(gzip.open(sys.argv[1])))
    elif sys.argv[1].endswith(".fa") or sys.argv[1].endswith(".txt"):
        input_fasta = file(sys.argv[1],'r')
    else:
        raise Exception("Unsupported File Type")
else:
    print """
    \tUsage:\n\t\tgenerate_masked_ranges.py <fasta file | .fa or .fa.gz> <chrome find> <chrome
replace>

    \t\t'Chrome find' and 'chrome replace' are used to find and replace the name of a chromsome. For
example,
    \t\treplacing CHROMSOME_I with chr1 can be accomplished by using the command as follows:
    \t\t\tpython generate_masked_ranges.py my_fasta.fa CHROMSOME_ chr
    \t\tOutput is to stdout
    """
    raise SystemExit


n, state = 0, 0 # line, character, state (0=Out of gap; 1=In Gap)
chrom, start, end = None, None, None

with input_fasta as f:
    for line in f:
        line = line.replace("\n","")
        if line.startswith(">"):
            # Print end range
            if state == 1:
```

```
            print '\t'.join([chrom ,str(start), str(n)])
            start, end, state  = 0, 0, 0
        n = 0 # Reset character
        chrom = line.split(" ")[0].replace(">","")
        # If user specifies, replace chromosome as well
        if len(sys.argv) > 2:
            chrom = chrom.replace(sys.argv[2],sys.argv[3])
    else:
        for char in line:
            if state == 0 and char == "N":
                state = 1
                start = n
            elif state == 1 and char != "N":
                state = 0
                end = n
                print '\t'.join([chrom ,str(start), str(end)])
            else:
                pass

            n += 1 # First base is 0 in bed format.

# Print mask close if on the last chromosome.
if state == 1:
        print '\t'.join([chrom ,str(start), str(n)])
        start, end, state  = 0, 0, 0
```

**Supplementary Note 2**. Snakefiles and associated code for SV calling pipeline

Snakefile1 main rules:

```
SAMPLES = ["Sample1", "Sample2"]

snakefiles = "snakefiles/"

include: snakefiles + "align"

include: snakefiles + "smoove"

rule all:

input:

"smoove/results/genotyped/paste/cohort.smoove.square.anno.vcf.gz",

"indexcov/index.html"
```

**Snakefile 1:**

```
rule bwa_index_reference:
input:

"/users/r01acb15/sharedscratch/reference/ICSASG_v2.fa"

output:

"/users/r01acb15/sharedscratch/reference/ICSASG_v2.fa"

log:

"bwa_index_reference.log"

shell:

"bwa index {input} &> {log}"

rule bwa_map:

input:

reference = "/users/r01acb15/sharedscratch/reference/ICSASG_v2.fa",

forward = "data/samples/{sample}_1.fastq",

reverse = "data/samples/{sample}_2.fastq"

output:

"mapped_reads/{sample}.bam"

params:
```

```
rg="@RG\\tID:{sample}\\tSM:{sample}\\tLB:lib1"

log:

"bwa_{sample}.log"

threads:

16

shell:

"(bwa mem -R '{params.rg}' -t {threads} {input.reference}

{input.forward} {input.reverse} | "

"samtools view -Sb - "

"> {output}) "

"2> {log}"

rule samtools_sort:

input:

"mapped_reads/{sample}.bam"

output:

"sorted_reads/{sample}.bam"

log:

"sort_{sample}.log"

shell:

"samtools sort -T sorted_reads/{wildcards.sample} "
"-O bam {input} > {output} "

"2> {log} "

rule samtools_index:

input:

"sorted_reads/{sample}.bam"

output:

"sorted_reads/{sample}.bam.bai"
```

shell:
"samtools index {input}"

rule goleft_indexcov:

input:

bam=expand("sorted_reads/{sample}.bam", sample=SAMPLES),

bai=expand("sorted_reads/{sample}.bam.bai", sample=SAMPLES)

output:

"indexcov/index.html"

log:

"indexcov.log"

shell:

"goleft indexcov -d indexcov/ {input.bam} 2> {log}"

**Snakefile 2:**

REF = "/users/r01acb15/sharedscratch/reference/ICSASG_v2.fa",
EXCLUDE =

"/users/r01acb15/sharedscratch/reference/high_depth_regions.min100_cluster.Nga
ps.bed",

GFF = "/users/r01acb15/sharedscratch/reference/Salmo_salar-annotation.gff3"
rule call:

input:

bam="sorted_reads/{sample}.bam"

output:

"smoove/results/{sample}-smoove.genotyped.vcf.gz"

conda:

"envs/smoove2.3.yaml"

shell:

"smoove call --outdir smoove/results/ --exclude {EXCLUDE} –name

{wildcards.sample} --fasta {REF} -p 1 --genotype {input.bam}"

rule merge:

input:

vcf=expand("smoove/results/{sample}-smoove.genotyped.vcf.gz", sample=SAMPLES)

output:

"smoove/results/merged.sites.vcf.gz"

conda:

"envs/smoove2.3.yaml"

shell:

"smoove merge --name merged -f {REF} --outdir ./smoove/results/

{input.vcf}"

rule genotype:

input:

merge="smoove/results/merged.sites.vcf.gz",

bam="sorted_reads/{sample}.bam"

output:

"smoove/results/genotyped/{sample}-joint-smoove.genotyped.vcf.gz"

conda:

"envs/smoove2.3.yaml"

shell:

"smoove genotype -d -x -p 1 --name {wildcards.sample}-joint –outdir smoove/results/genotyped/ --fasta {REF} --vcf {input.merge} {input.bam}"

rule paste:

input:

vcf=expand("smoove/results/genotyped/{sample}-jointsmoove.

genotyped.vcf.gz", sample=SAMPLES)

output:

"smoove/results/genotyped/paste/cohort.vcf.gz"

conda:

"envs/smoove2.3.yaml"

shell:
"smoove paste --name cohort {input.vcf}"

rule annotate:

input:

"smoove/results/genotyped/paste/cohort.vcf.gz"

output:

"smoove/results/genotyped/paste/cohort.smoove.square.anno.vcf.gz"

conda:

"envs/smoove2.3.yaml"

shell:

"smoove annotate --gff {GFF} {input} | bgzip -c > {output}"

**Conda environment to run smoove step of Snakefile:**

This environment should be inserted into a .yaml file and executre with conda.

name: smoove2.3

channels:

- bioconda
- conda-forge
- defaults

dependencies:

- anaconda=custom=py27h4a00acb_0
- asn1crypto=0.24.0=py27_3
- backports=1.0=py_2
- backports.functools_lru_cache=1.5=py_1
- backports_abc=0.5=py_1
- bcftools=1.9=h4da6232_0
- bedtools=2.27.1=he941832_2
- blas=1.0=mkl
- boto3=1.9.4=py_0
- botocore=1.12.4=py_0
- bzip2=1.0.6=h470a237_2
- ca-certificates=2019.6.16=hecc5488_0
- certifi=2019.6.16=py27_0
- cffi=1.11.5=py27h5e8e0c9_1
- cryptography=2.3.1=py27hdffb7b8_0
- cryptography-vectors=2.3.1=py27_0
- curl=7.61.0=h93b3f91_2
- cycler=0.10.0=py_1
- cytoolz=0.9.0.1=py27h470a237_1

- dbus=1.13.0=h3a4f0e9_0
- docutils=0.14=py27_1
- duphold=0.0.9=0
- enum34=1.1.6=py27_1
- expat=2.2.5=hfc679d8_2
- fontconfig=2.13.1=h65d0f4c_0
- freetype=2.9.1=h6debe1e_4
- functools32=3.2.3.2=py27_2
- futures=3.2.0=py27_0
- gawk=4.2.1=h470a237_0
- gettext=0.19.8.1=h5e8e0c9_1
- glib=2.55.0=h464dc38_2
- gsort=0.0.6=1
- gst-plugins-base=1.12.5=hde13a9d_0
- gstreamer=1.12.5=h61a6719_0
- htslib=1.9=hc238db4_4
- icu=58.2=hfc679d8_0
- idna=2.7=py27_2
- intel-openmp=2019.0=117
- ipaddress=1.0.22=py_1

**Supplementary Note 3**: Custom R script used to obtain Fst values from random comparisons and establish probability value for outlier SVs

```
#######200 iterations to obtain Fst values from random comparisons######

###"Indivduals.txt" is a file with the IDs of the animals in the farmed and South Norway populations. One ID per line.

###"TOTAL2mb_highqual.vcf" is VCF file with the calls for the high quality SVs

#We calculate Fst values for 200 random subsets of 34 and 257 animals

for i in {1..200}
do
  shuf -n 34 individuals.txt > small_pop.txt

##Obtain 34 random animals
  grep -w -v -F -f small_pop.txt individuals.txt > large_pop.txt

##Select the remaining 257 animals
  vcftools --vcf TOTAL2mb_highqual.vcf --weir-fst-pop small_pop.txt --weir-fst-pop large_pop.txt --out fst_pop1_vs_pop2               ##Perform Fst calculation using VCFtools
  cat fst_pop1_vs_pop2.weir.fst >> fst_results_n200.txt

##Save the results of all the iterations to the file "fst_results_n200.txt"
done


#########P-values for each individual high quality SV########

###"Farmed-vs-SouthNorwayWild.fst" is a file with the VCFtools Fst calculation output for the farmed vs wild South Norway comparison for the high-quality SVs. One SV per line, first and second columns have the position of the SNP, the third column has the Fst value.
cat Farmed-vs-SouthNorwayWild.fst | while read line

##We process one SV each time

do

FST=$(echo $line | awk '{print $3;}')

#We extract the Fst value for the current SV

echo $line | awk '{print $1,$2;}' | sed 's/ /\t/g' - | grep -w -f - fst_results_n200.txt | cut -f3 - > temporal.txt

#From the previous 200 random Fst iterations, we select the lines for the current SV

OVER=$(awk -v var="$FST" '$1>var{c++} END{print c+0}' temporal.txt)

#We obtain the number of random iterations that resulted in a higher Fst than the one obtain in random vs wild South Norway for the current SNP

echo $OVER >> number_below_fst.txt
```

#We save the number of higher random Fst in a file, the number for each SV in one line
PVAL=$(bc <<< "scale=4;$OVER/200")

#We calculate the proportion of higher random Fst by dividing the number by 200. This is the p-value
echo

$PVAL >> p_value.txt

#We store the p-values in a file, one line per SV
done

paste Farmed-vs-SouthNorwayWild.fst number_below_fst.txt p_value.txt > Farmed-vs-SouthNorwayWild-with-pval.fst

#We merge the output of VCFtools Fst calculation with the p-values
